# Mathematical characterization of population dynamics in breast cancer cells treated with doxorubicin

**DOI:** 10.1101/2021.12.01.470781

**Authors:** Emily Y. Yang, Grant R. Howard, Amy Brock, Thomas E. Yankeelov, Guillermo Lorenzo

## Abstract

The development of chemoresistance remains a significant cause of treatment failure in breast cancer. We posit that a mathematical understanding of chemoresistance could assist in developing successful treatment strategies. Towards that end, we have developed a model that describes the effects of the standard chemotherapeutic drug doxorubicin on the MCF-7 breast cancer cell line. We assume that the tumor is composed of two subpopulations: drug-resistant cells, which continue proliferating after treatment, and drug-sensitive cells, which gradually transition from proliferating to treatment-induced death. The model is fit to experimental data including variations in drug concentration, inter-treatment interval, and number of doses. Our model recapitulates tumor growth dynamics in all these scenarios (as quantified by the concordance correlation coefficient, CCC > 0.95). In particular, superior tumor control is observed with higher doxorubicin concentrations, shorter inter-treatment intervals, and a higher number of doses (*p* < 0.05). Longer inter-treatment intervals require adapting the model parameterization after each doxorubicin dose, suggesting the promotion of chemoresistance. Additionally, we propose promising empirical formulas to describe the variation of model parameters as functions of doxorubicin concentration (CCC > 0.78). Thus, we conclude that our mathematical model could deepen our understanding of the effects of doxorubicin and could be used to explore practical drug regimens achieving optimal tumor control.

## Introduction

Breast cancer is the most common cancer among women worldwide and the leading cause of cancer death in over 100 countries [1]. Chemotherapy is a primary component of cancer treatment and options have both advanced and increased considerably in recent years [2]. However, the development of chemoresistance, and resulting tumor recurrence, remains a common cause of treatment failure and a primary cause of cancer death [3],[4]. Indeed, for a standard chemotherapy drug such as doxorubicin, chemoresistance can develop within just 6-10 months [5],[6].

From a biological perspective, the development of chemoresistance is governed by many complex mechanisms, such as treatment-induced genetic and epigenetic alterations, altered metabolic states, and adaptive responses of the tumor microenvironment [7]–[10]. Tumor cells can also possess an intrinsic phenotypic or genetic resistance that can render the therapy ineffective even before acquired chemoresistance develops [11]–[13]. Moreover, the existence of intratumoral heterogeneity and its role in tumor regrowth have become increasingly recognized, as the presence of even a minor subpopulation of drug-resistant cells can give rise to tumor relapse [9],[14]–[16]. Furthermore, phenotype switching, in which tumor cells swap between varying degrees of drug-resistant and drug-sensitive phenotypes, can enable the establishment of more permanent chemoresistance mechanisms that hinder complete tumor eradication [9],[17]–[19]. In light of the complex biological processes underlying chemoresistance development, we believe that a robust framework is needed to comprehensively integrate the growing knowledge of this phenomenon and guide future research efforts. To this end, we propose that experimentally-validated mathematical models of chemoresistance mechanisms could be a potent tool in understanding the dynamics of overall tumor drug response. The description of cancer growth and therapeutic response by leveraging mechanistic mathematical models is a rich field known as mathematical oncology [20]–[22]. This approach has already shown promise in characterizing breast cancer growth and treatment response in both the preclinical and clinical settings [23]–[30].

There are several mechanistic approaches to mathematically describe chemoresistance [31], with the original theoretical models dating back more than two decades [32],[33]. The standard strategy consists of defining a multicompartmental tumor cell population including one or multiple species of both drug-resistant and drug-sensitive cells, which evolve and interact over time following a set of ordinary differential equations, or over both space and time according to a set of partial differential equations [34]– [38]. Alternatively, Sun *et al*. utilized a stochastic, multiscale model that incorporated heterogeneous population dynamics with drug pharmacokinetics and microenvironment contributions to drug resistance in melanoma patients [39]. Furthermore, Pisco *et al*. and Álvarez-Arenas *et al*. applied the evolutionary theories of Darwinian selection and Lamarckian induction to guide their modeling of drug resistance in leukemia cells and non-small cell lung carcinoma, respectively [40],[41]. However, despite these promising studies, there is a still a dearth of experimentally-validated mechanistic models of chemoresistance in breast cancer, with which we could test alternative biological hypotheses to ultimately enhance chemotherapeutic strategies for individual patients. For instance, Chapman *et al*. developed a model integrating phenotypic switching of cell differentiation states and tumor heterogeneity to characterize therapeutic escape in the triple-negative subtype, but the empirical validation of their model predictions currently remains limited [42]. Additionally, *in vitro* studies usually label cell lines as homogeneously drug-resistant or drug-sensitive and assume a static drug sensitivity [43],[44], which overlooks the existence of intratumoral heterogeneity and transient drug resistance. Moreover, preclinical studies often assess tumor cell death at a single time point 24-72 hours post-treatment [45]–[47]. This experimental setting does not enable the characterization of long-term tumor drug responses and, hence, the development of drug-induced chemoresistance.

Here, we present a mechanistic model to describe the dynamics of drug response and chemoresistance development in MCF-7 breast cancer cells treated with doxorubicin, which we fit to time-resolved microscopy measurements of tumor cell number subjected to diverse therapeutic plans over long experimental times (>8 days). Doxorubicin is an anthracycline drug that is extensively used in chemotherapeutic regimens for breast cancer [30],[48],[49] and whose mechanism of action induces tumor cell death [50]–[52]. Our work continues the first efforts of Howard *et al*. in studying doxorubicin resistance in breast cancer cell populations by leveraging several experimentally-informed mechanistic models [34],[53]. While Howard *et al*. originally proposed multiple models to characterize this phenomenon and selected the best of them for each dataset, we have developed a single model that can be extended for multiple drug doses and is also amenable to an adaptive parametrization with each doxorubicin dose. To incorporate intratumoral heterogeneity, we assume that the doxorubicin treatment induces a compartmentalization of the breast cancer cell population into two subgroups: surviving cells and cells that will die due to doxorubicin action. This compartmentalization ultimately results from the underlying distribution of diverse drug sensitivity phenotypes in the breast cancer cell population. The surviving cells, which we classify as resistant with respect to the treatment, continue proliferating after exposure to doxorubicin, while the remainder of the cells, which we classify as sensitive to the treatment, progressively transition from proliferation to drug-induced death. Hence, drug sensitivity is assumed to be dynamic with time, thereby accounting for tumor cell plasticity [9],[17]–[19]. Additionally, our model is fit to the same time-resolved microscopy experiments used in Howard *et al*. [53], in which breast cancer cells were subjected to doxorubicin treatments varying in either drug concentration, inter-treatment interval, or the number of doses. Our results show that our proposed model can fit the data observed in all three scenarios with remarkable accuracy. We have also analyzed the model parameter trends for each experiment and built empirical parameter formulas as functions of doxorubicin concentration, which may provide further insight into the development of chemoresistance.

The remainder of this work is organized as follows. First, we describe the procedures to acquire and process our time-resolved microscopy data. We also describe the derivation of the model and explain the numerical and statistical methods leveraged in this study. We then present the results from our model fittings for each of the three aforementioned experimental scenarios and analyze the corresponding quality of fit and trends in model parameters. To conclude, we discuss the main implications from our work, its limitations, and future directions.

## Methods

### Data acquisition

As complete data acquisition procedures are provided elsewhere [34], here we present only the salient details.

#### Cell culture

MCF-7 human breast cancer cells (ATCC HTB-22) were cultured in Minimum Essential Media (Gibco) supplemented with 10% fetal bovine serum (Gibco) and 1% penicillin-streptomycin (Gibco). Cells were maintained at 37° C with 5% CO_2_. A stable fluorescent cell line expressing constitutive EGFP with a nuclear localization signal (MCF7-EGFPNLS1) was established to aid in the automated cell quantification of the time resolved microscopy measurements [34],[53]. Genomic integration of the EGFP expression cassette was accomplished by leveraging the Sleeping Beauty transposon system. The EGFP-NLS sequence was obtained as a gBlock (IDT) and cloned into the optimized Sleeping Beauty transfer vector pSBbi-Neo (which was a gift from Eric Kowarz, Addgene plasmid #60525) [54]. To mediate genomic integration, this two-plasmid system consisting of the transfer vector containing the EGFP-NLS sequence and the pCMV(CAT)T7-SB100 plasmid containing the Sleeping Beauty transposase was co-transfected into the MCF-7 population utilizing Lipofectamine 2000 (mCMV(CAT)T7-SB100 was a gift from Zsuzsanna Izsvak, Addgene plasmid #34879)[55]. After gene integration with Sleeping Beauty transposase, EGFP+ cells were collected by fluorescence activated cell sorting and maintained in media supplemented with 200 ng/mL G418 (Caisson Labs) in place of penicillin-streptomycin.

#### Doxorubicin response experiments

Cells were seeded in a 96-well plate at a target density of 2,000 cells/well and grown for approximately 48 hours to allow for cell adhesion and recovery from passaging. An IncuCyte S2 Live Cell Analysis System (Essen/Sartorius, Goettingen, Germany) was used to collect fluorescent and phase contrast images every 2-4 hours. Images were collected for periods of 21-56 days to ensure that cultures in which cells recover after exposure to doxorubicin, were able to display logistic growth. Doxorubicin treatment was prepared by reconstituting doxorubicin hydrochloride (Cayman Chemical 15007, Ann Arbor, Michigan) in water and mixing it with 100 µL of growth media at 2× the target concentration, which was then added to each well of the plate. The drug-containing media was then replaced with fresh growth media after 24 hours. Three experiment types were run, in which either the doxorubicin concentration, the inter-treatment interval, or the number of doses was varied (see Table 1). Each doxorubicin concentration was tested in n = 6 replicates, while each inter-treatment interval and number of doses was tested in n = 12 replicates.

**Table 1.** Experimental conditions. In Experiment 1, one dose of doxorubicin was delivered at concentrations varying from 10 to 300 nM (n = 6). In Experiment 2, two doses of 75 nM doxorubicin were delivered at inter-treatment intervals varying from 0 to 16 days (n = 12). In Experiment 3, one to five doses of 75 nM doxorubicin were delivered at either 2-day or 2-week inter-treatment intervals (n = 12).

| Experiment | Concentration [nM] | Inter-treatment Interval [d] | Number of Doses | Number of Replicates |
| --- | --- | --- | --- | --- |
| 1 | 0, 10, 20, 35, 50, 75, 100, 125, 150, 300 | - | 1 | 6 |
| 2 | 75 | 0, 2, 4, 6, 8, 10, 12, 14, 16 | 2 | 12 |
| 3 | 75 | 2 or 14 | 1, 2, 3, 4, 5 | 12 |

### Data preprocessing

#### Image Analysis

Using IncuCyte’s integrated software, cell quantification was performed on the fluorescent images using the green fluorescence channel. Individual cells were consistently resolved using standard image analysis techniques of background subtraction, followed by thresholding, edge detection, and minimum area filtering. The phase contrast images were consulted in parallel to aid the validation of image analysis [34],[53].

#### Data truncation

The time courses extracted from some wells did not provide meaningful data throughout the entire time course due to a variety of reasons. These included the cell population growing to confluence and fluctuating with feeding cycles, being disturbed during media replenishment, or growing three-dimensionally resulting in cells overlapping each other and thereby compromising the ability to accurately quantify cell numbers. Thus, for each dataset, the estimated cell number was truncated either just prior to reaching confluence, when the cell number dropped more than 50% due to media handling, or when repeated discontinuities were observed in the time course data.

#### Data normalization

For smaller discontinuities in which less than 50% of the cells were lost, the data was normalized by dividing the cell number at time points before the discontinuity by a constant α [34],[53], calculated *via* Eq. (1):

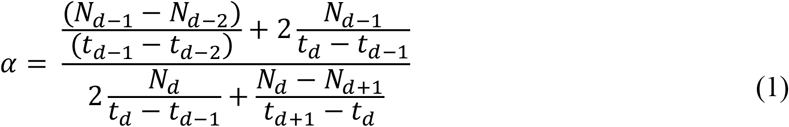

in which *N*_*d*_, *N*_*d-i*_, and *N*_*d+i*_ are the cell numbers at the discontinuity, *i* points before the discontinuity, and *I* points after the discontinuity, respectively. *t*_*d*_, *t*_*d-i*_, and *t*_*d+i*_ are the times of the discontinuity, *i* points before the discontinuity, and *i* points after the discontinuity, respectively. The objective of this normalization was to smooth the first and second derivatives of the cell number across the discontinuity [34],[53].

#### Outlier removal

For replicates that possessed outliers, the *rmoutliers* function from MATLAB R2020b (The Mathworks, Natick, MA) was used to remove data points using median filtering. A visual inspection of the resulting data confirmed that this method removed evident outliers from the original series, while maintaining the natural fluctuations in tumor cell counts (see Supplementary Figures S1-S5).

### Mathematical model

We present a mathematical model to describe the response of MCF-7 breast cancer cells to treatment with doxorubicin in the three experimental scenarios listed in Table 1. We begin by describing the biological mechanisms captured by the model assuming a single dose of doxorubicin (Experiment 1, Table 1). Then, we show how the model can be generalized to multiple doses (Experiments 2 and 3, Table 1), and can also be modified to vary specific parameters with each dose. Figure 1 illustrates the main tumor cell dynamics described by our model after each dose of doxorubicin, which are further detailed in the following paragraphs. The reader can refer to Supplementary Table S1 for a consolidated list of model parameter definitions and their units.

**Figure 1.**
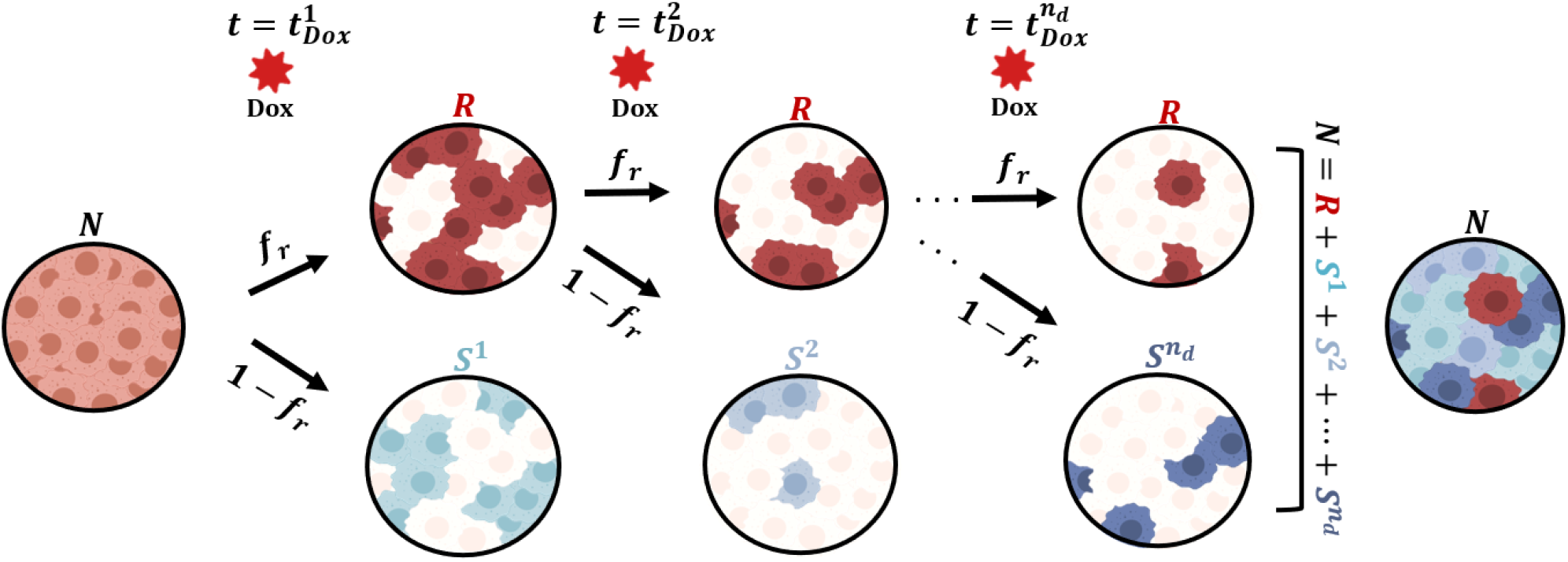
Generalized model of tumor cell response to multiple doses of doxorubicin treatment. We start with a population of untreated tumor cells and let them grow for approximately 48 hours. At time 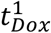, we add a dose of doxorubicin (Dox) to each well. We assume that after the treatment, the tumor cells exhibit either one of two responses: drug-resistant (*R*) or drug-sensitive (*S*^1^). The fraction of cells in either subpopulation is determined by *f*_*r*_, the fraction of drug-resistant cells. After the subsequent doses (*i* = 1,2,…,*n*_*d*_), we assume that a fraction *f*_*r*_ of the drug-resistant cells survive the treatment, while a fraction (1 - *f*_*r*_) induces a new subpopulation of sensitive cells (*S*^*i*^), such that the total number of tumor cells is 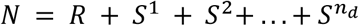 for times 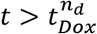. This Figure was created using BioRender.com.

#### Single-dose model

We start with a population of tumor cells (*N*) that grow untreated for a specified period of time prior to doxorubicin treatment (approximately 48h). We assume that these untreated cells follow logistic growth:

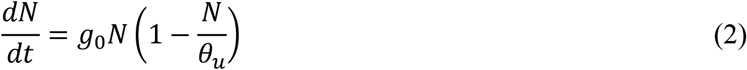

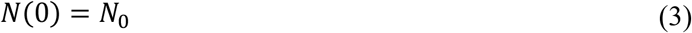

where *g*_*0*_ is the untreated proliferation rate, θ_*u*_ the untreated tumor cell carrying capacity, and *N*_0_ is the initial number of tumor cells. We set θ_*u*_ = 53,873 cells, which corresponds to the mean value resulting from the fitting of Eqs. (2-3) to the untreated datasets in Experiment 1 (i.e., 0 nM doxorubicin; further details can be found in Supplementary Tables S2-S5 and Supplementary Figure S1).

Let 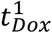 denote the time at which a single dose of doxorubicin is delivered, as described in Experiment 1 (Table 1). At this time point, we assume that a fraction *f*_*r*_ of the tumor cells exhibits a drug-resistant response (*R*), while a fraction 1 - *f*_*r*_ exhibits a drug-sensitive response (*S*). We denote the initial number of drug-resistant and drug-sensitive cells right after the treatment with doxorubicin as 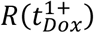 and 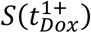, respectively, which are defined based on the number of untreated cells immediately before the delivery of doxorubicin, 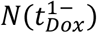, as

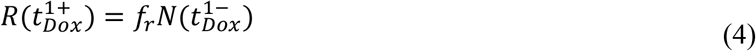

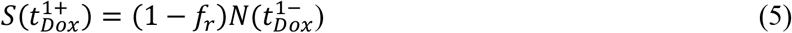

such that the total tumor cell number *N* for time 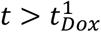 is calculated as

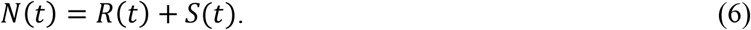

Note that Eqs. (4)-(6) ensure the continuity in the tumor cell count before and after the treatment with doxorubicin, as observed in the corresponding experimental data (see Supplementary Figure S2).

We assume that the drug-resistant cells also follow logistic growth with a different rate and carrying capacity:

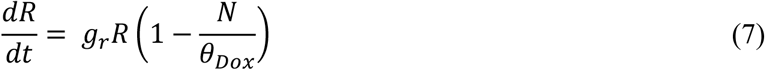

where *g*_*r*_ is the proliferation rate of drug-resistant tumor cells and θ_*Dox*_ is the treated tumor cell carrying capacity. For the drug-sensitive cells, we assume that their logistic growth dynamics gradually transition from proliferation to treatment-induced death at an exponentially-decaying rate:

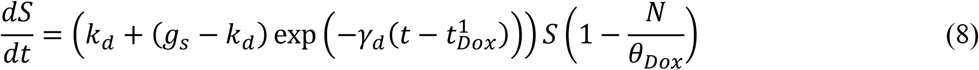

where *k*_*d*_ and *g*_*s*_ denote the drug-induced death rate and the proliferation rate of drug-sensitive tumor cells, respectively, while γ_*d*_ represents the drug-induced death delay rate.

To estimate the treated tumor cell carrying capacity (θ_*Dox*_) in Eqs. (7)-(8), we used either one of two approaches. If the last tumor cell count in a dataset was greater than 30% of θ_*Dox*_, then was fit along with the other model parameters. Conversely, if the last tumor cell count was less than 30% of θ_*u*_ then we fixed θ_*Dox*_ to the mean of the values obtained from the replicates of the same experiment in which this parameter was directly fit. The rationale for this approach is that we observed that final tumor cell counts below 30% of θ_*u*_ did not provide enough identifiability for θ_*Dox*_, which ultimately induced significant model fitting errors.

#### Multiple-dose model

Let us now consider a treatment schedule consisting of *n*_*d*_ doses of doxorubicin delivered at times 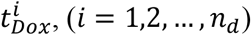, (*i* = 1,2,…,*n*_*d*_), as described in Experiments 2 and 3 (Table 1). For the first dose, the multiple-dose model remains identical to the single-dose model described in the previous section. For the subsequent drug doses, we assume that a fraction *f*_*r*_ of the drug-resistant cells before treatment 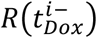, continue to exhibit this response, while a fraction 1 - *f*_*r*_ gives rise to a new drug-sensitive subpopulation *S*^*i*^. Thus, after the delivery of the *i*^*th*^ dose (*i* ≥2, the number of drug-resistant tumor cells 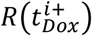 and the initial number of the new subpopulation of drug-sensitive cells 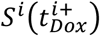 are calculated as

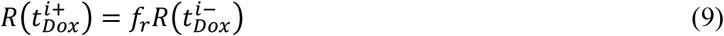

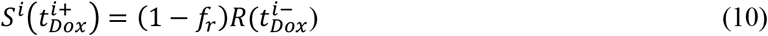

such that the total number of tumor cells during and after treatment with multiple doses of doxorubicin is given by

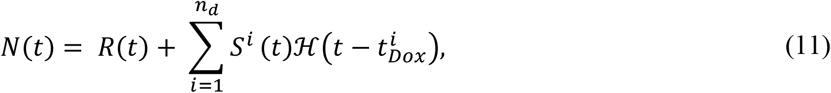

where 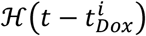 is the Heaviside step function, which equals 0 for 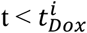 and 1 for 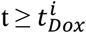. Note that Eqs. (8)-(11) ensure the continuity in the total tumor cell number before and after each doxorubicin dose, as observed in the data from Experiments 2 and 3 (Table 1; see Supplementary Figures S3-S5).

In this multiple-dose model, we assume that the drug-resistant cells continue to follow logistic growth after each of the consecutive doxorubicin doses, as described by Eq. (7). Additionally, each of the *i*^*th*^ drug-sensitive subpopulations is assumed to follow the growth dynamics defined in Eq. (8). Thus, for *i* = 1,2,…,*n*_*d*_, the dynamics of each drug-sensitive subpopulation *S*^*i*^ is given by

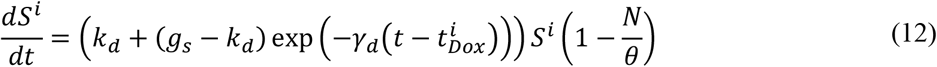

Finally, we consider two versions of the multiple-dose model: one featuring constant parameters, and another in which we vary *f*_*r*_ and γ_*d*_ with the delivery of each dose. Our underlying hypothesis is that longer inter-treatment intervals require an adaptive parameterization because they contribute to the development of chemoresistance [34], [56]–[58], which would be represented in our model by higher fractions of resistant cells (*f*_*r*_) along with drug-sensitive subpopulations (*S*^*i*^) exhibiting longer transition times from proliferation to treatment-induced death (i.e., lower values of γ_*d*_). In the Results section, we show that these hypotheses are significantly supported by the fitting of these two model versions to the data from Experiments 2 and 3 (Table 1).

### Numerical methods

#### Model fitting

We fit the single-dose model to the time-course data from Experiment 1 (Table 1), and we fit the multiple-dose model to the time-course data from Experiments 2 and 3 (Table 1). Model fitting was carried out with a nonlinear least-squares method, *via* the MATLAB (R2020b) function *lsqnonlin*. We leveraged a trust-region reflective algorithm with function, step, and optimality tolerances of 10^−6^, while the maximum number of function evaluations and iterations was set to 20,000. The parameter bounds and initial guesses were guided by the results from Howard *et al*. [34], and are summarized in Supplementary Tables S2, S8, S12, S16 and S17. The ordinary differential equations in our models were solved using a Runge-Kutta method as provided by *ode45* in MATLAB (R2020b).

#### Empirical parameter formulas

We constructed empirical formulas for the single-dose model parameters as a function of doxorubicin concentration based on the model fittings to the datasets from Experiment 1 (Table 1). To this end, we also applied a nonlinear least-squares method using a trust-region reflective algorithm provided by *lsqnonlin* in MATLAB (R2020b), as described in the previous section. The initial guess and bounds for the empirical parameters in these formulas were chosen according to the range of the single-dose model parameter values obtained from the fittings to the datasets from Experiment 1 (Table 1). The medians of the distributions of these fitted model parameters at each doxorubicin concentration were used as the observed values for the empirical parameter formula fits. Based on the observed trends of the fitted single-dose model parameters, we chose different empirical equations to describe their change as a function of doxorubicin concentration (e.g., an exponential decay; see the Results section and Supplementary Tables S6 and S7 for further details).

### Statistical analysis

To assess our model’s quality of fit to the time course data, we calculated the coefficient of determination (*R*^2^), the normalized root mean square error (NRMSE), the Pearson correlation coefficient (PCC), and the concordance correlation coefficient (CCC) [59]. In the Results section, we report the median and range of these metrics across all the replicates of each experiment. More detailed values can be found in the Supplementary Tables S4, S10, S14, S20. Nonlinear regression parameter confidence intervals and nonlinear regression prediction confidence intervals were calculated using *nlparci* and *nlpredci* in MATLAB (R2020b), respectively. To test for significant differences between two values of a model parameter or quality-of-fit metric within each experimental scenario, we performed two-sided Wilcoxon rank-sum tests with 5% significance using *ranksum* in MATLAB (R2020b).

To assess the validity of the proposed empirical formulas using the single-dose model fittings, we ran a simulation test in which we qualitatively compared the model outcomes based on these formulas with the corresponding experimental observations at each drug concentration. To this end, Latin hypercube sampling based on *lhsdesign* in MATLAB (R2020b) was used to define 200 parameter combinations assuming uniform distributions over the 95% confidence intervals of the fitted empirical parameter formulas at each doxorubicin concentration, as calculated by *nlpredci* in MATLAB (R2020b).

## Results

### Fitting the single-dose model to Experiment 1 data: varying doxorubicin concentrations

Figure 2 shows representative model fits for the observed growth of MCF-7 cell populations treated with only one dose of doxorubicin at concentrations ranging from 10 to 300 nM (Experiment 1, Table 1). Model fits for all replicates at each drug concentration (n = 6) can be found in Supplementary Figure S2. We report the median and range of all the fitted model parameters for each doxorubicin concentration in Supplementary Table S3, while Figure 3 shows the boxplots of the fitted parameter distributions for each doxorubicin concentration. The median and range of the quality of fit metrics for the single-dose model fits to Experiment 1 data were: NRMSE (3.33 [0.80, 12.43]), *R*^2^ (>0.99 [0.96, >0.99]), PCC (>0.99 [0.98, >0.99]), and CCC (0.99 [0.97, >0.99]). Supplementary Table S4 further provides detailed quality of fit metrics for each doxorubicin concentration. Figure 2, Supplementary Figure S2, and Supplementary Table S3 show that, as doxorubicin concentration is increased, the drug-resistant cells exhibit a decrease in growth rate and number, while the drug-sensitive cells undergo a faster transition from proliferation to treatment-induced death. These trends ultimately lead to significantly lower final total tumor cell counts (< 0.05, see Supplementary Table S5) and larger delay or even suppression of tumor regrowth in the cells exposed to higher doxorubicin concentrations (see Supplementary Figure S2), suggesting that tumor control improves as the doxorubicin dose is increased.

**Figure 2.**
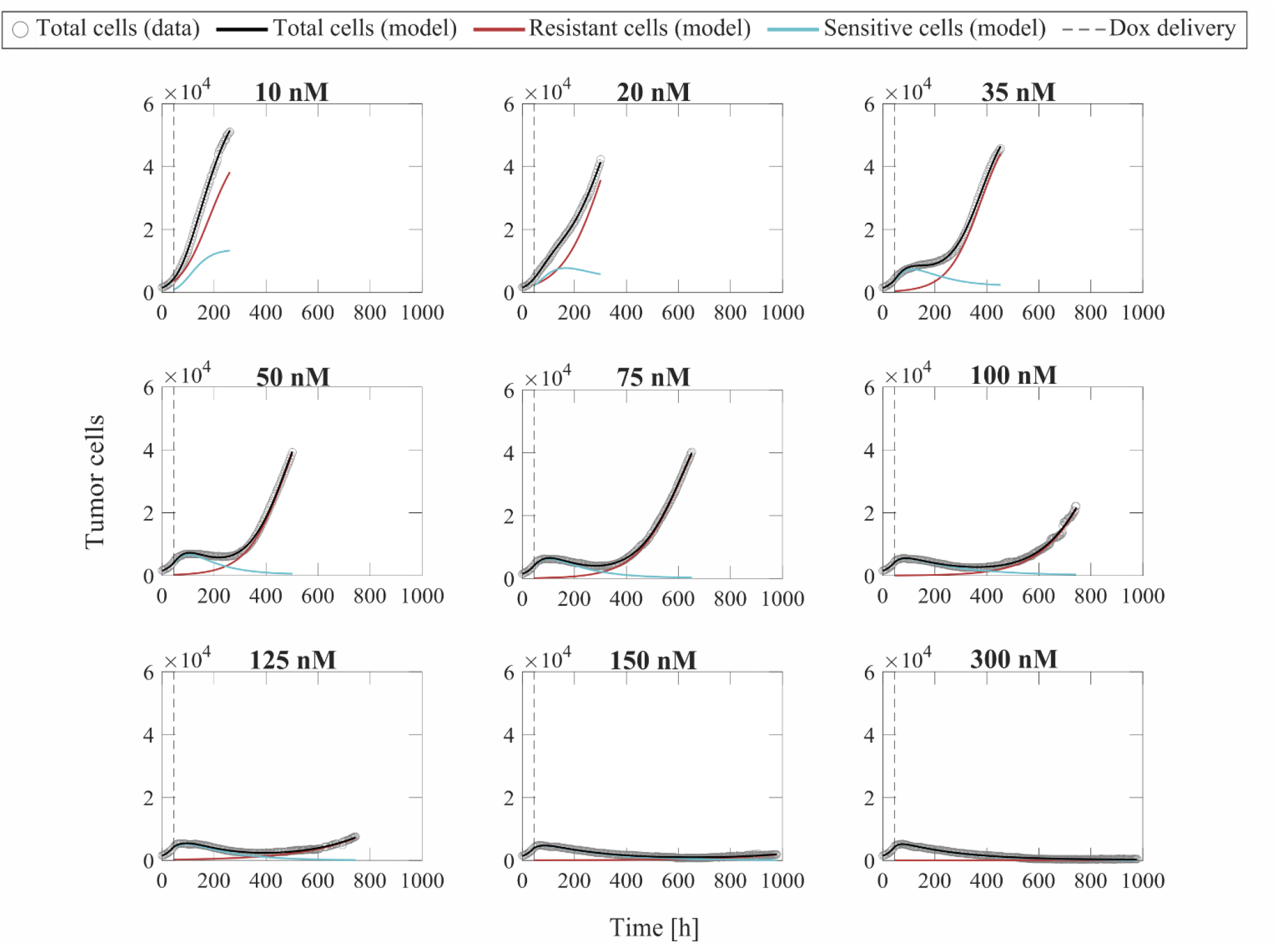
Representative fits of the single-dose model for varying concentrations of doxorubicin. Data and model fittings are shown for a representative replicate treated with 10 to 300 nM doxorubicin concentrations (Experiment 1, Table 1). Experimental data are shown in gray circles. The number of total cells, resistant cells, and sensitive cells obtained with the fitted single-dose model are shown in black, red, and blue solid lines, respectively. The time of doxorubicin delivery is represented with a vertical grey dashed line. As doxorubicin concentration is increased, we observe a decrease in the growth rate of drug-resistant cells and a faster transition from growth to treatment-induced death in the drug-sensitive cells. These drug-induced effects ultimately translate into a longer delay (or even suppression) of tumor growth post-treatment and lower total tumor cell count for higher doxorubicin concentrations, indicating superior tumor control overall. The median and range of the quality of fit metrics across all replicates in Experiment 1 (Table 1) are NRMSE: 3.33 [0.80, 12.43], *R*^2^ : >0.99 [0.96, >0.99], PCC: >0.99 [0.98, >0.99], and CCC: 0.99 [0.97, >0.99].

**Figure 3.**
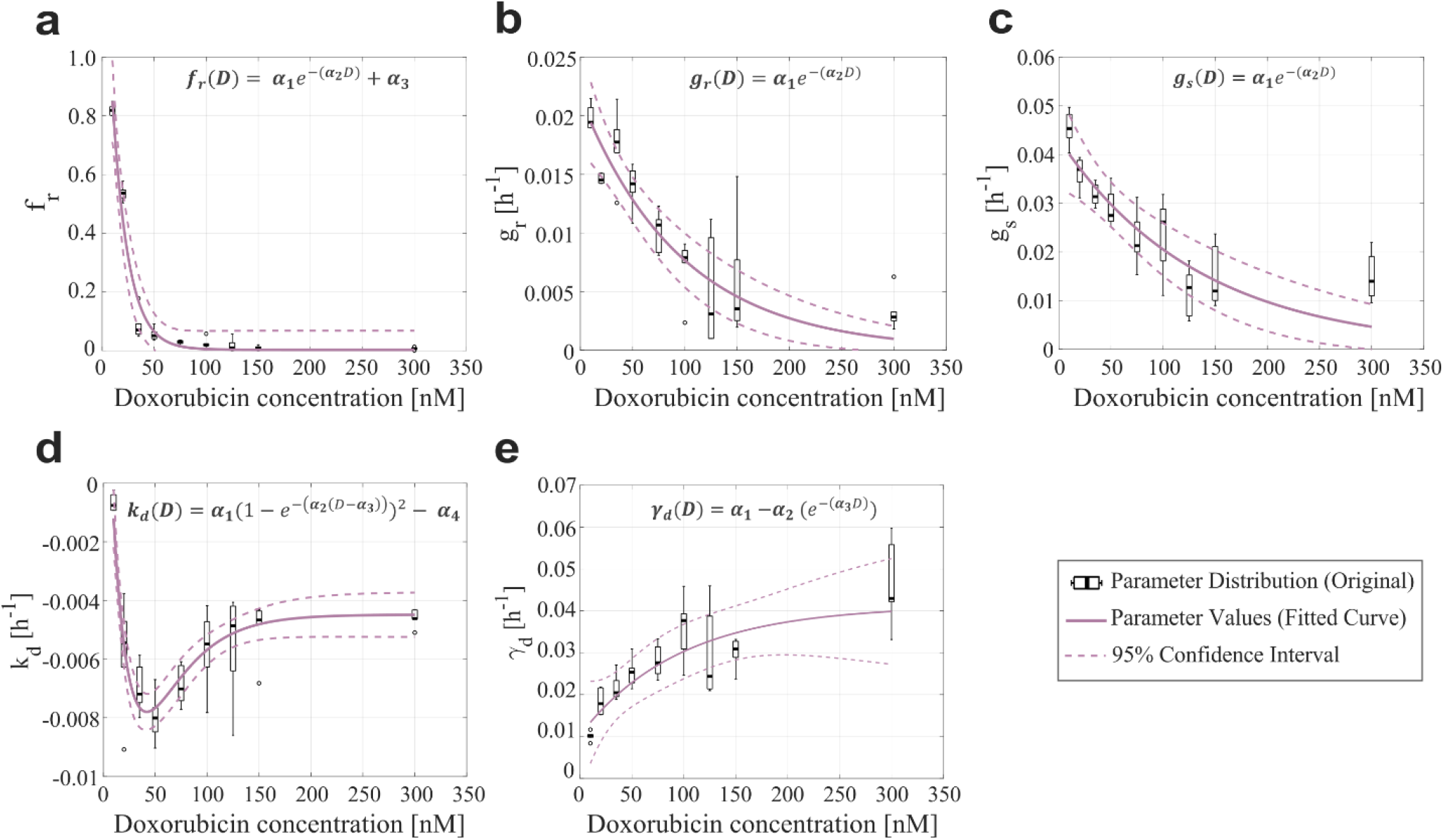
Empirical parameter formulas for varying doxorubicin concentrations. The proposed empirical formulas indicated at the top of each panel (a-e) were fit to the median of the corresponding parameter distributions obtained from fitting the single-dose model to the varying concentration datasets from Experiment 1 (Table 1). *D* denotes doxorubicin concentration in nM, while α_*i*_ (*i* = 1,2,…) are empirical parameters. The distributions of the single-dose model parameters are represented with black boxplots, in which outliers are represented as black circles. The resulting curves from fitting the empirical parameter formulas are shown as purple solid lines, and their corresponding 95% confidence intervals are plotted as purple dashed lines. Panel (a) shows the parameter formula for the fraction of resistant cells (*f*_*r*_). Panel (b) shows the parameter formula for the proliferation rate of drug-resistant tumor cells (*g*_*r*_). Panel (c) shows the parameter formula for the proliferation rate of drug-sensitive tumor cells (*g*_*s*_). In panels (a) – (c), we observe that as the drug concentration increases, the corresponding single-dose model parameter values decrease exponentially. Panel (d) shows the parameter formula for the doxorubicin-induced death rate in drug-sensitive cells (*k*_*d*_), which we approximated with an equation based on a Morse-potential relationship. Panel (e) shows the parameter formula for the doxorubicin-induced death delay rate of drug-sensitive cells (γ_*d*_), which increases and then plateaus as the drug concentration increases. Median and range of quality of fit metrics are NRMSE: 17.65 [5.45, 153.6], *R*^2^ : 0.91 [0.73, 0.98], PCC: 0.95 [0.86, 0.99], and CCC: 0.85 [0.76, 0.88].

Figure 3 shows the fitted empirical formulas for the fraction of resistant cells (*f*_*r*_), the drug-resistant cell proliferation rate (*g*_*r*_), the drug-sensitive cell proliferation rate (*g*_*s*_), the doxorubicin-induced death rate of drug-sensitive cells (*k*_*d*_), and the doxorubicin-induced death delay rate of drug-sensitive cells (γ_*d*_). These empirical formulas are functions of doxorubicin concentration, which is denoted with *D*. The fitted empirical parameter values and their confidence intervals can be found in Supplementary Table S6, while the corresponding quality of fit metrics can be found in Supplementary Table S7. For *f*_*r*_, *g*_*r*_ and *g*_*s*_ we observe a clear exponentially decaying trend as drug concentration is increased (Fig. 3a-c). In the case of *f*_*r*_, we added an additional constant empirical parameter to the decaying exponential to ensure that the empirical formula captures the low nonzero values of this parameter for the higher doxorubicin concentrations (otherwise, the exponential decay would reach the horizontal asymptote at *f*_*r*_ = 0 for low doxorubicin concentrations). The parameter *k*_*d*_ exhibits a complex trend, consisting of a steep decreasing branch for doxorubicin concentrations under 50 nM, followed by an increasing branch that plateaus for doxorubicin concentrations over 150 nM. We found that an empirical formula based on a Morse-potential relationship [60] captured this trend (Fig. 3d). For γ_*d*_, we chose a decaying exponential flipped with respect to the horizontal axis to capture the increasing trend that ultimately plateaus at a nonzero value (Fig. 3e). The median and range of the quality of fit metrics for the proposed empirical formulas were: NRMSE (17.65 [5.45, 153.6]), *R*^2^ (0.91 [0.73, 0.98]), PCC (0.95 [0.86, 0.99]), and CCC (0.85 [0.76, 0.88]). We note that the NRMSE of the fitted empirical formula for *f*_*r*_ reached values beyond 100%. This is due to the small values of *f*_*r*_ at high concentrations of doxorubicin, where the NRMSE is not relevant to modeling outcomes; for reference, the RMSE for the fitted empirical formula for *f*_*r*_ is 0.0476.

Once the parameter formulas had been established, we wanted to qualitatively assess the range of tumor growth dynamics that our formulas could reproduce. For each doxorubicin concentration, we sampled the 95% confidence intervals of the fitted empirical parameter formulas (dashed purple lines in Fig. 3) using Latin hypercube sampling to obtain 200 parameter combinations, with which we ran corresponding model simulations. Figure 4 presents the median and range of the model simulations plotted against the median and range of the experimental data for each doxorubicin concentration tested in Experiment 1 (Table 1). We observe that the proposed empirical parameter formulas (Fig. 3) are able to predict a wide range of model solutions and that the simulations are able to capture the overall tumor growth dynamics observed in the datasets from Experiment 1 (Table 1).

**Figure 4.**
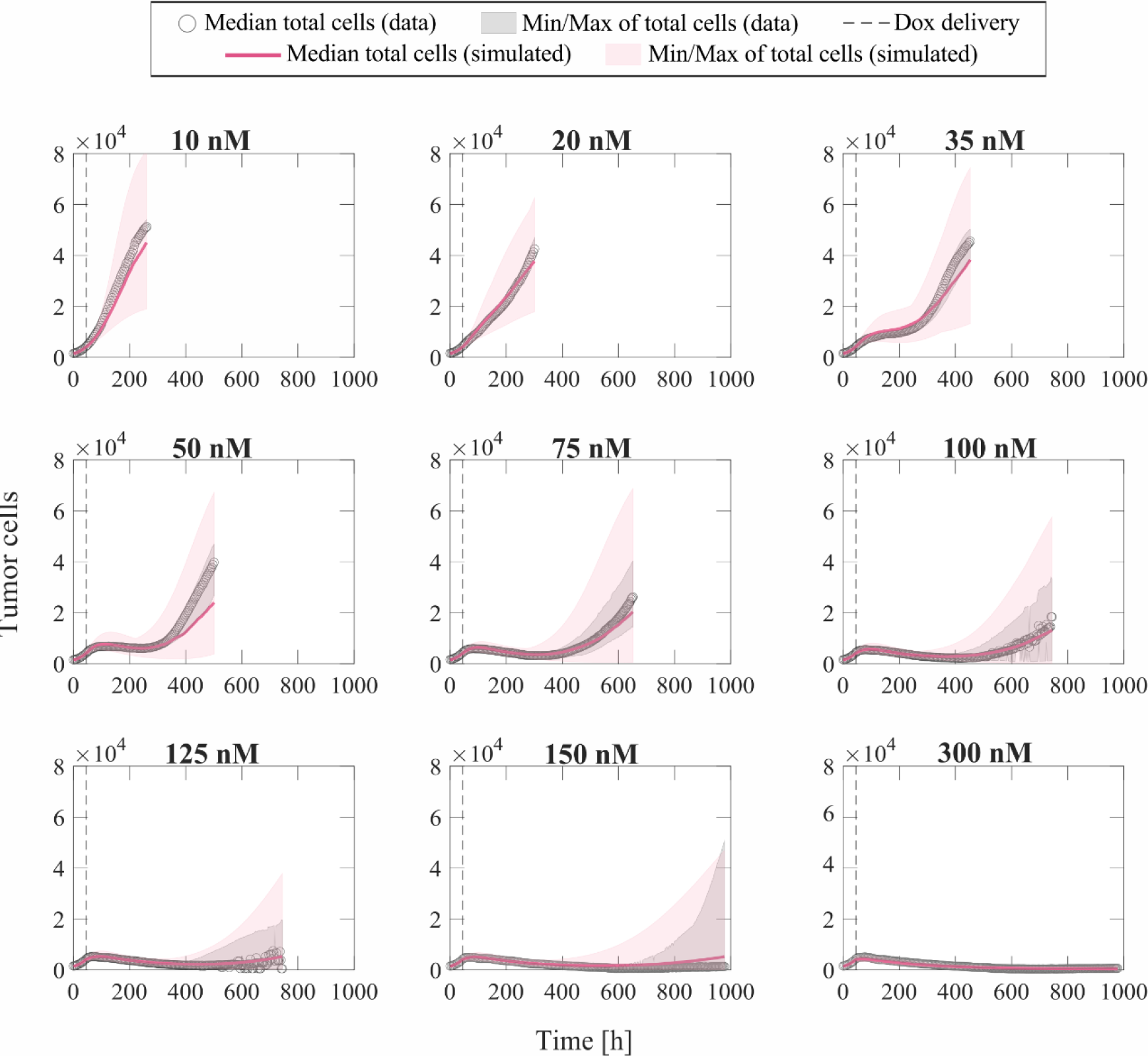
Comparison of simulated tumor growth based on empirical parameter formulas with respect to experimental data for varying doxorubicin concentrations. We sampled the 95% confidence intervals for the fitted empirical parameter formulas in Figure 3 using Latin hypercube sampling to obtain 200 parameter combinations for each doxorubicin concentration, with which we carried out corresponding simulations with the single-dose model. The median and range of the model simulations are plotted with the median and range of the experimental data from Experiment 1 (Table 1) for comparison. The median of the experimental data is shown with gray circles, and the range of the experimental data is represented with gray shaded regions. The median of the model simulations is plotted as a pink solid line, and the range of the simulations is shown as pink shaded regions. The time of doxorubicin delivery is represented with a vertical grey dashed line. We observe that our fitted parameter formulas from Figure 3 can reproduce a wide range of tumor cell dynamics, including the tumor cell growth observed in the varying concentration datasets (Experiment 1, Table 1).

### Fitting the multiple-dose model to Experiment 2 data: varying inter-treatment intervals

To fit the experimental data for varying inter-treatment intervals (Experiment 2, Table 1), we initially used the two versions of the multiple-dose model; i.e., with all parameters held constant or varying *f*_*r*_ and γ_*d*_ with each drug dose. Figure 5 shows the distribution of the NRMSE in fitting the datasets at each inter-treatment interval (n = 12) for both models. We observe a significant difference between the NRMSEs obtained with either version of the multiple-dose model, such that the varying and model provides a significantly lower NRMSE at 8-, 10-, 12-, 14-, and 16-day inter-treatment intervals (*p* : 0.0043, 0.0017, 2.46 × 10^−4^, 5.92 × 10^−4^, 3.66 × 10^−4^, respectively). Furthermore, the model with constant parameters provides a significantly lower NRMSE at the 2-day inter-treatment interval (*p* = 0.0024). Thus, the results shown in Figure 5 justify the use of the model with constant parameters for inter-treatment intervals shorter than 8 days and the model with varying *f*_*r*_ and γ_*d*_ for inter-treatment intervals ≥ 8 days. We followed this model selection criterion for fitting the datasets from Experiments 2 and 3 (Table 1) for the remainder of this work.

**Figure 5.**
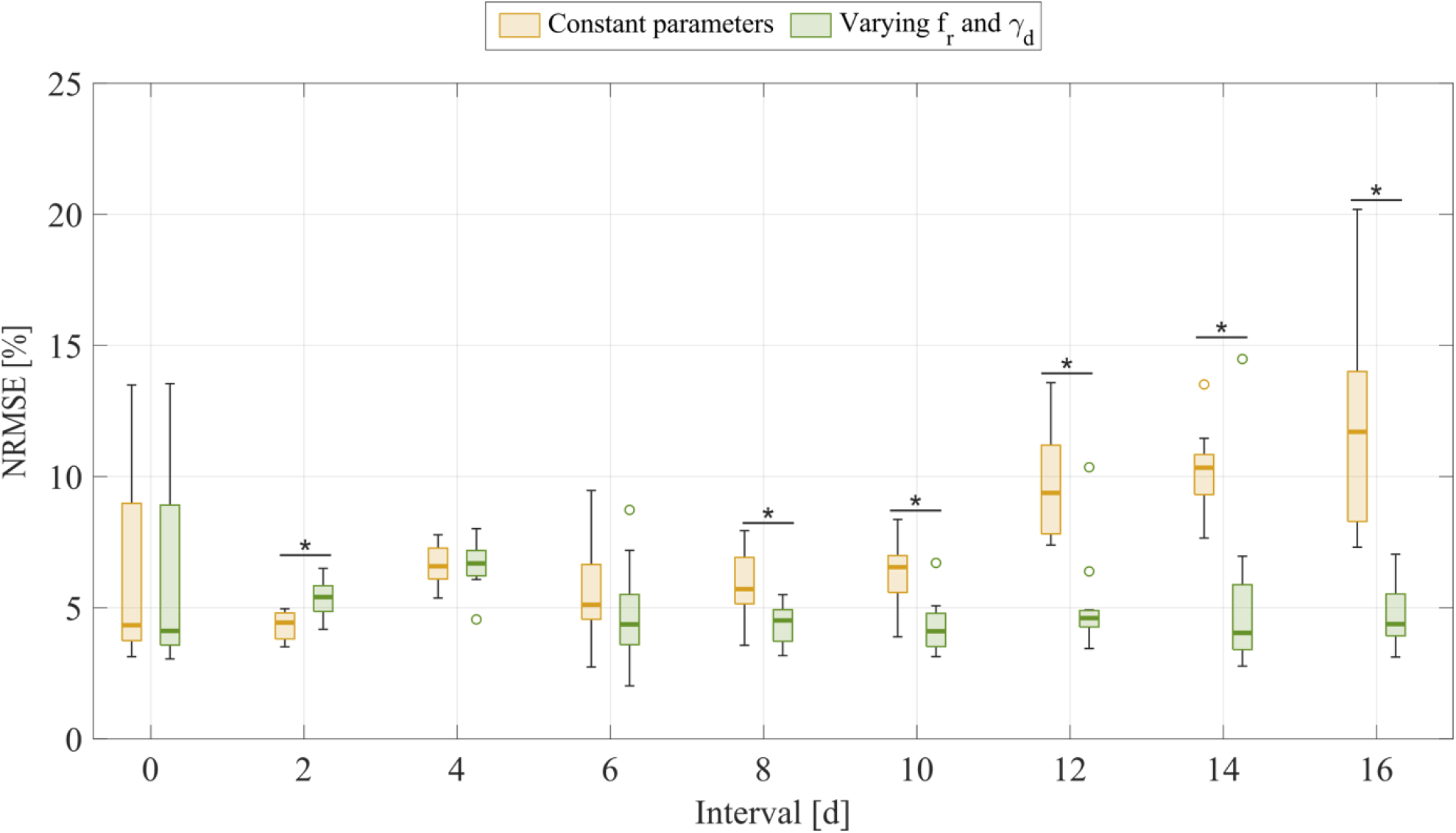
Comparison of fitting the experimental data for varying inter-treatment intervals with the multiple-dose model with constant versus varying parameters. For each inter-treatment interval tested in Experiment 2 (Table 1), we compared the normalized root mean squared error (NRMSE) calculated from the fittings using the multiple-dose model with constant parameters (yellow boxplots) with the NRMSE calculated from the fittings using the multiple-dose model with varying and (green boxplots). Outliers are represented with circles. At inter-treatment intervals of 8, 10, 12, 14, and 16 days, there is a significantly lower NRMSE when the model with varying *f*_*r*_ and γ_*d*_ is used (*p* : 0.0043, 0.0017, 2.46, 5.92, 3.66, respectively). Additionally, we observe that, with an inter-treatment interval of 2 days, there is a significantly lower NRMSE when the model with constant parameters is used (*p* 0.0024). An asterisk (^*^) indicates *p* <0.05 (two-sided Wilcoxon rank sum test).

Figure 6 shows representative model fits for the observed growth of MCF-7 cell populations treated with two doses of 75 nM doxorubicin delivered at inter-treatment intervals ranging from 0 to 16 days (Experiment 2, Table 1). Model fits for all the replicates at each inter-treatment interval (n = 12) can be found in Supplementary Figure S3. Additionally, Supplementary Table S9 summarizes the median and range of the fitted model parameters for each inter-treatment interval. The median and range of the quality of fit metrics were: NRMSE (4.64 [2.74, 14.3]), *R*^2^ (0.99 [0.80, >0.99]), PCC (>0.99 [0.91, >0.99]), and CCC (0.99 [0.90, >0.99]). More detailed quality of fit metrics for each inter-treatment interval are reported in Supplementary Table S10. As the inter-treatment interval lengthens, the drug-resistant cells tend to adopt an increasingly larger growth rate and the drug-sensitive cells transition more slowly from proliferation to drug-induced death after two doses of doxorubicin treatment (see Figure 6, Supplementary Figure S3, and Supplementary Table S9). These effects appear to promote tumor regrowth after the second dose in most replicates for inter-treatment intervals of 6 days or longer and after the first dose for inter-treatment intervals of 12 days or longer. Overall, this ultimately leads to significantly higher final total tumor cell counts as the inter-treatment interval is lengthened (*p* < 0.05, see Supplementary Table S11), suggesting that increased time spans between consecutive doses of doxorubicin is conducive to poorer tumor control.

**Figure 6.**
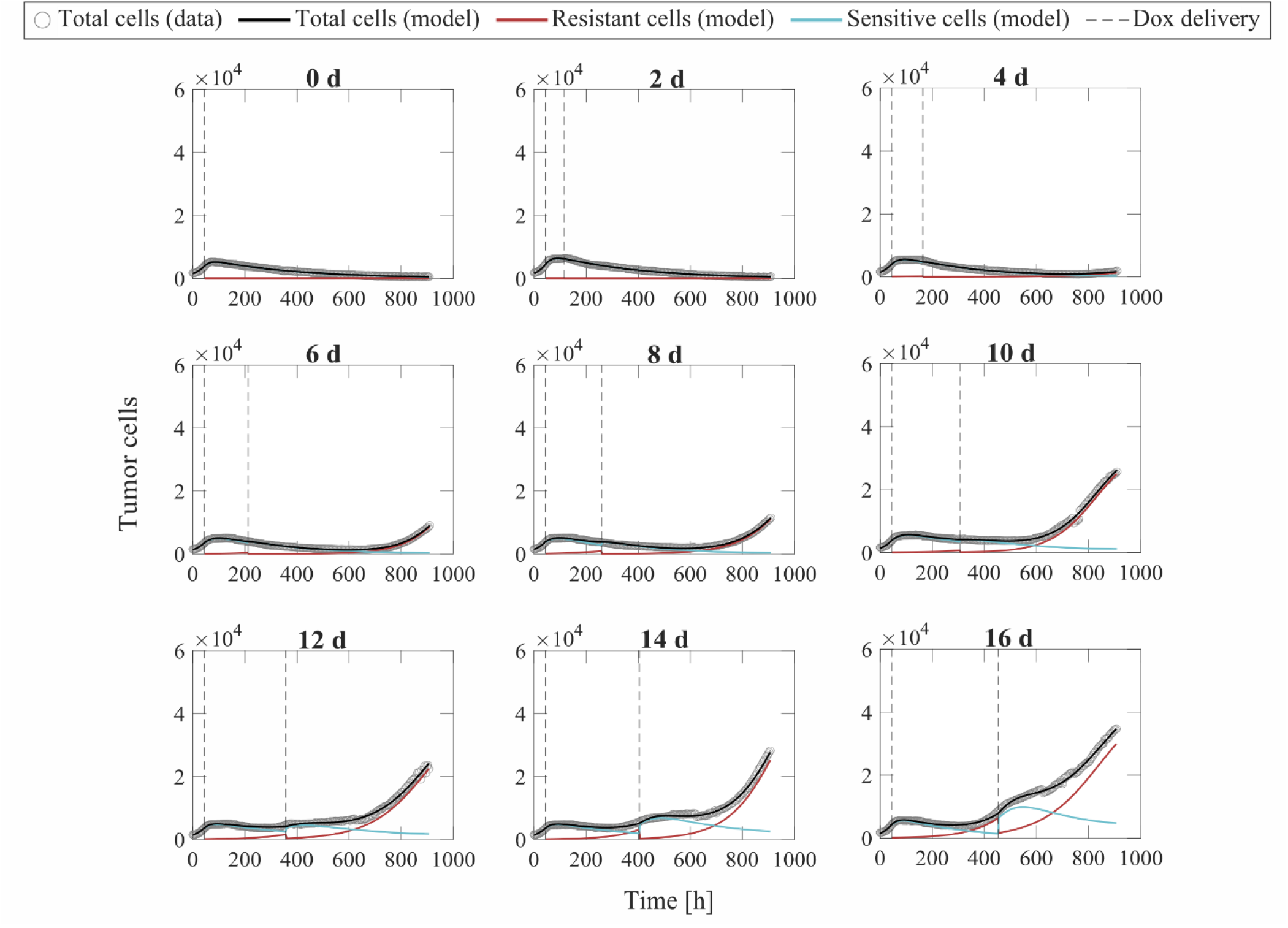
Representative fits of the multiple-dose model for varying inter-treatment intervals. Data and model fittings are shown for a representative replicate exposed to two doses of 75 nM doxorubicin delivered at inter-treatment intervals ranging from 0 to 16 days (Experiment 2, Table 1). Experimental data are shown in gray circles. The number of total cells, resistant cells, and sensitive cells obtained with the fitted multiple-dose model are shown in black, red, and blue solid lines, respectively. The times of doxorubicin delivery are represented with vertical grey dashed lines. For inter-treatment intervals of 0 to 6 days, the multiple-dose model with constant parameters was used for data fitting. For inter-treatment intervals of 8 to 16 days, we used the multiple-dose model with varying *f*_*r*_ and γ_*d*_. As the inter-treatment interval is lengthened, we observe an increase in the growth rate of drug-resistant cells and a slower transition from growth to treatment-induced death in the drug-sensitive cells. These drug-induced effects ultimately lead to a tumor relapse after the second dose for inter-treatment intervals of 6 days or longer in most replicates, as well as tumor regrowth after the first dose for inter-treatment intervals of 12 days or longer. These observations suggest increasingly poor tumor control as the two doses of 75 nM of doxorubicin are spaced further out in time. The median and range of the quality of fit metrics across all datasets in Experiment 2 (Table 1) are NRMSE: 4.64 [2.74, 14.3], *R*^2^ : 0.99 [0.80, >0.99], PCC: >0.99 [0.91, >0.99], and CCC: 0.99 [0.90, >0.99].

Additionally, Figure 7 shows the distributions of the fitted *f*_*r*_ and γ_*d*_ values from fitting the multiple-dose model to the data with varying inter-treatment intervals (Experiment 2, Table 1). When the model with constant parameters is used (inter-treatment intervals from 0 to 6 days), we observe a trend towards higher resistant fractions and delayed transitions to treatment-induced death in drug-sensitive cells as the two doses are further spaced in time. This observation is further supported by the distributions of varying *f*_*r*_ and γ_*d*_ obtained from fitting the multiple-dose model to the data for inter-treatment intervals from 8 to 16 days. After the second dose, the resistant fraction significantly increases and the transition from proliferation to treatment-induced death in drug-sensitive cells significantly slows (*p* < 0.05, see Fig. 7), thereby suggesting an enhanced chemoresistance in both tumor cell subpopulations for longer inter-treatment intervals. Moreover, the distributions shown in Figure 7 further support the use of the multiple-dose model with varying *f*_*r*_ and γ_*d*_ for longer inter-treatment intervals.

**Figure 7.**
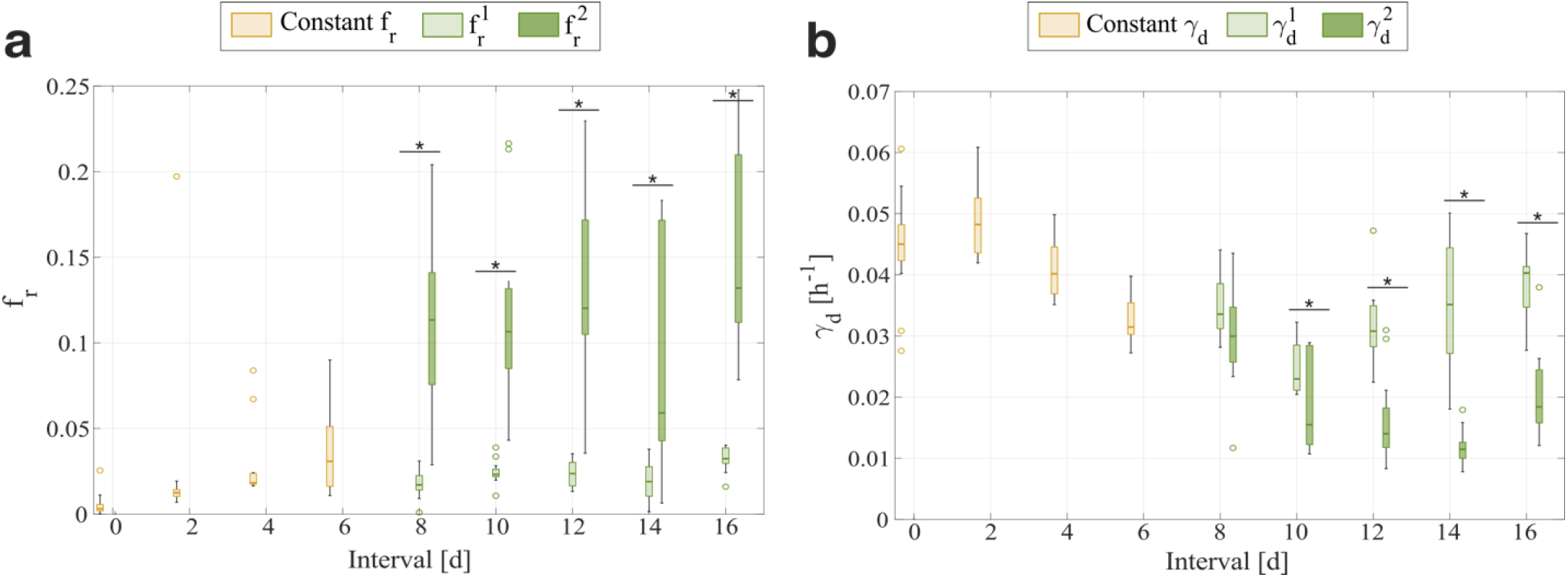
Comparison of the *f*_*r*_ and γ_*d*_ distributions obtained from fitting the multiple-dose model to the experimental data for varying inter-treatment intervals. The parameter distributions are represented as boxplots and were obtained from fitting the multiple-dose model to the varying inter-treatment interval datasets from Experiment 2 (Table 1). Outliers are represented with circles. Panel (a) shows the distributions for the fraction of resistant cells (*f*_*r*_). Panel (b) shows the distributions for the doxorubicin-induced death delay rate in drug-sensitive tumor cells (γ_*d*_). For 0 to 6 day inter-treatment intervals, *f*_*r*_ and γ_*d*_ are kept constant in the model (yellow boxplots); whereas, for 8 to 16 day inter-treatment intervals, we vary *f*_*r*_ and γ_*d*_ with each doxorubicin dose (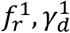 : light green boxplots, 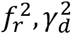 : dark green boxplots). As the inter-treatment interval is lengthened from 0 to 6 days, the constant *f*_*r*_ and γ_*d*_ show a trend towards higher resistant fractions and slower transitions to doxorubicin-induced death, suggesting increasingly poorer tumor control. When *f*_*r*_ and γ_*d*_ are varied with each dose, we observe that the second *f*_*r*_ values correspond to significantly higher resistant fractions for 8-, 10-, 12-, 14-, and 16-day inter-treatment intervals (*p* : 4.7 × 10^−5^, 3.7 × 10^−5^, 3.7 × 10^−5^, 3.7 × 10^−5^, and 3.7 × 10^−5^, respectively) and that the second γ_*d*_ values represent significantly slower transitions to treatment-induced death for 10-, 12-, 14-, and 16-day inter-treatment intervals (*p* : 0.0141, 3.7 × 10^−5^, 6.0 × 10^−5^, and 9.7 × 10^−5^ respectively). These changes in *f*_*r*_ and γ_*d*_ after the second dose also suggest an increasingly poorer tumor control after the second dose with a longer inter-treatment interval. An asterisk (^*^) indicates *p* <0.05 (two-sided Wilcoxon rank sum test).

### Fitting the multiple-dose model to Experiment 3 data: varying number of doses

Figure 8 shows representative model fits for the observed growth of MCF-7 cell populations treated with 1 to 5 doses of 75 nM doxorubicin delivered at either 2-day or 2-week inter-treatment intervals (Experiment 3, Table 1). The datasets from the cells treated with a 2-day inter-treatment interval were fitted with the multiple-dose model with constant parameters, while the datasets from the cells treated with a 2-week inter-treatment interval were fitted with the multiple-dose model with varying *f*_*r*_ and γ_*d*_. Model fits for all the replicates for each number of doses and both inter-treatment intervals (n = 12) can be found in Supplementary Figures S4 and S5. The median and range of the fitted model parameters for each dose number are summarized in Supplementary Tables S13, S18, and S19. For the replicates treated every 2 days, the median and range of the quality of fit metrics were: NRMSE (12.2 [2.72, 19.1]), *R*^2^ (0.99 [0.87, >0.99]), PCC (0.99 [0.93, >0.99]), and CCC (0.99 [0.93, >0.99]). Likewise, for the replicates treated every 2 weeks, the median and range of the quality of fit metrics were: NRMSE (3.21 [1.91, 8.57]), *R*^2^ (>0.99 [0.93, >0.99]), PCC (>0.99 [0.97, >0.99]), CCC (0.99 [0.96, >0.99]). More detailed quality of fit metrics for each number of doses and both inter-treatment intervals can be found in Supplementary Tables S14 and S20.

**Figure 8.**
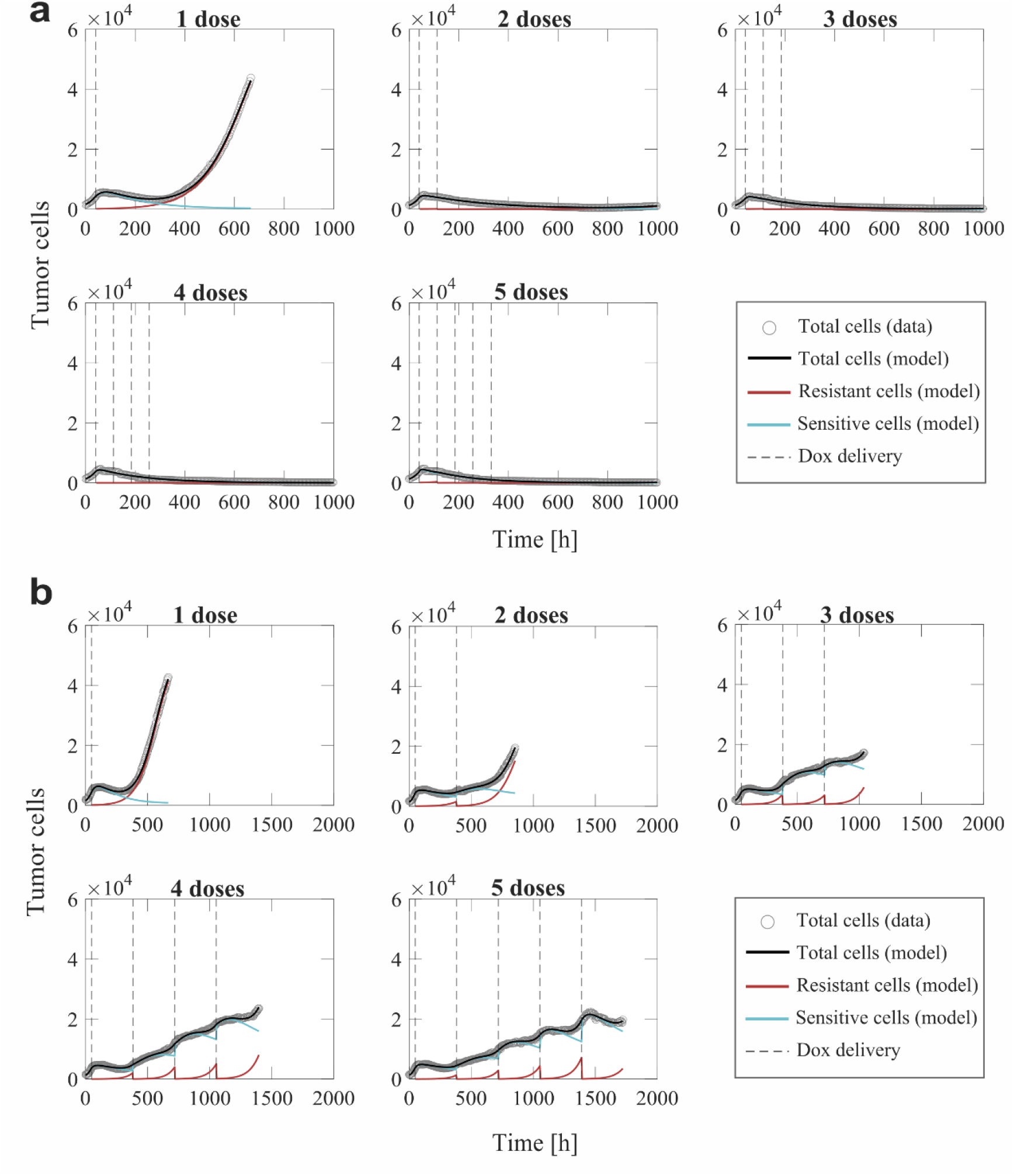
Representative fits of the multiple-dose model for a varying number of doxorubicin doses. Data and model results are shown for a representative replicate treated with 1 to 5 doses of 75 nM doxorubicin delivered at either 2-day or 2-week inter-treatment intervals (Experiment 3, Table 1). Experimental data are shown in gray circles. The number of total cells, resistant cells, and sensitive cells obtained with the fitted multiple-dose model are shown in black, red, and blue solid lines, respectively. The times at which doxorubicin is delivered are represented with vertical grey dashed lines. Panel (a) shows fittings for 1 to 5 doxorubicin doses delivered at 2-day inter-treatment intervals obtained with the model with constant parameters. The median and range of the quality of fit metrics across all replicates for this Experiment 3 subgroup (Table 1) are NRMSE: 12.2 [2.72,19.1], *R*^2^ : 0.99 [0.87,>0.99], PCC: 0.99 [0.93,>0.99], CCC: 0.99 [0.93,>0.99]. Panel (b) shows fittings for 1 to 5 doxorubicin doses delivered at 2-week inter-treatment intervals obtained with the model with varying *f*_*r*_ and γ_*d*_. The median and range of the quality of fit metrics across all replicates for this Experiment 3 subgroup (Table 1) are NRMSE: 3.21 [1.91,8.57], *R*^2^ : >0.99 [0.93,>0.99], PCC: >0.99 [0.97, >0.99], CCC: 0.99 [0.96,>0.99]. Overall, we observe that there is superior tumor control with an increased number of doses, which is further improved when the doses are delivered at shorter inter-treatment intervals. As the inter-treatment interval is lengthened from 2 days to 2 weeks, we observe that the growth rate and number of the drug-resistant cells increase, while the drug-sensitive cells exhibit a slower transition from proliferation to treatment-induced death.

The model fittings plotted in Figure 8 and Supplementary Figures S4 and S5 show that increasing the number of doses contributed to improved tumor control for the two inter-treatment intervals investigated in this work. In general, for the cells treated every 2 days, we observed significantly lower final total tumor cell counts as the number of doses was increased (*p* < 0.05, see Supplementary Table S15). Furthermore, delivering two or more doses effectively suppressed tumor growth at the end of the experiment, typically showing a decreasing branch in the total tumor cell count right after the first dose. When the inter-treatment interval was extended to 2 weeks, delivering more than one dose of doxorubicin also contributed to limited tumor cell growth (< 0.05, see Supplementary Table S21); however, most of the replicates showed an increasing trend in total tumor cell count over the experiment duration. Thus, with a 2-week inter-treatment interval, an increasing number of doses can decelerate tumor cell growth, but it cannot suppress it as observed with a 2-day inter-treatment interval. Furthermore, the model fitting results reported in Figure 8, Supplementary Figure S4 and S5, and Supplementary Tables S13, S18, and S19 show that, as the inter-treatment interval is lengthened from 2 days to 2 weeks, the drug-resistant cells exhibit a larger growth rate, while the drug-sensitive cells emerging after the second and subsequent doses undergo a slower transition to treatment-induced death. These effects, induced by the lengthened inter-treatment interval, contribute to explaining the superior tumor control in the 2-day experiments and align with the corresponding results shown in Figure 6, Supplementary Figure S3, and Supplementary Table S9.

We further investigated tumor cell dynamics for the Experiment 3 data with 2-week inter-treatment intervals by analyzing the evolving distributions of parameters *f*_*r*_ and γ_*d*_, which are shown in Figure 9. We observe that the drug-resistant fraction corresponding to the 1^st^ to 4^th^ doses 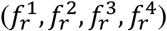 shows an increasing trend, which is indicative of progressive chemoresistance during treatment and aligns with the corresponding results shown in Figure 7. However, the fitted values for 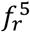 are significantly lower than the value obtained for 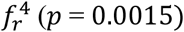. Additionally, we observe that the values for 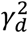 are significantly lower than that of 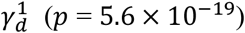, following the trend observed in Figure 7 for the data from Experiment 2. However, the values for 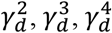, and 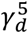 exhibit an increasing trend, with 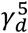 being significantly larger than 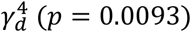. These changes in *f*_*r*_ and γ_*d*_ suggest that delivering multiple doses of doxorubicin may progressively limit or even revert the chemoresistance observed in the initial drug-resistant and drug-sensitive subpopulations.

**Figure 9.**
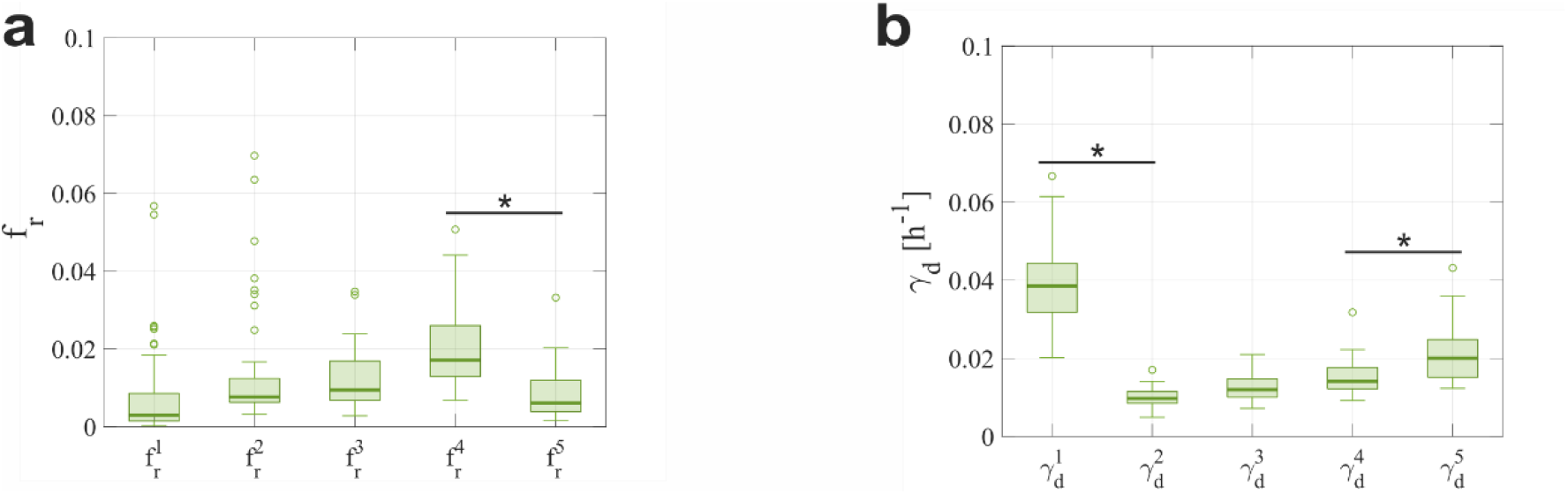
Distributions of *f*_*r*_ and γ_*d*_ obtained from fitting the multiple-dose model to the experimental data for a varying number of doses with a 2-week inter-treatment interval. The parameter distributions are represented as boxplots and were obtained from fitting the multiple-dose model to the 2-week inter-treatment interval datasets from Experiment 3 (Table 1), in which the model with varying *f*_*r*_ and γ_*d*_ was used. Outliers are represented as circles. Panel (a) shows the distributions of the fraction of resistant cells, such that a new value for *f*_*r*_ is defined for each drug dose (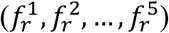). We observe an increasing trend in the first four *f*_*r*_ parameters, which suggests an increasing chemoresistance with each dose. However, 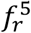 takes on significantly lower values than 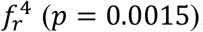, which suggests that adding more doses may limit the trend towards chemoresistance. Panel (b) shows the distributions of the doxorubicin-induced death delay rate of drug-sensitive cells, such that a new value of γ_*d*_ is defined for each drug dose 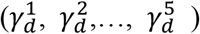. The values for 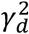 are significantly lower than those of 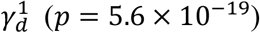. Hence, the second sensitive subpopulation shows a slower transition to treatment-induced death. However, the subsequent doxorubicin doses induce drug-sensitive subpopulations exhibiting an increasing γ_*d*_, with 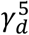 being significantly larger than 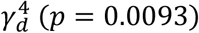. This observation further suggests that past a certain number of doses, initial chemoresistance appears to be reverted. An asterisk (^*^) indicates *p*<0.05 (two-sided Wilcoxon rank sum test).

## Discussion

We have presented a mathematical framework to describe the therapeutic response of MCF-7 breast cancer cells to treatment with doxorubicin and the development of chemoresistance in the *in vitro* setting. Our mathematical models feature drug-resistant/drug-sensitive tumor cell response and dynamic drug sensitivity to capture the underlying phenotypic heterogeneity and tumor plasticity, respectively. We presented a single-dose model that can be extended to a multiple-dose model, in which parameterization can vary with each dose. We fitted our models to various time-resolved microscopy datasets, which enabled us to evaluate tumor growth dynamics with our models in three experimental scenarios that varied either the doxorubicin concentration, the inter-treatment interval, or the number of doses (see Table 1). In all three cases, our models recapitulated the experimental observations, achieving a remarkable quality of fit.

In Experiment 1 (Table 1), we evaluated the effect of a single dose of doxorubicin on MCF-7 breast cancer cell growth and we found that tumor control was significantly improved with increased drug concentration (*p* < 0.05, see Supplementary Table S5). Our single-dose model showed that, at a subpopulation level, these dynamics emerged from a lower growth rate of drug-resistant cells and a faster transition from proliferation to treatment-induced death in drug-sensitive cells. The dynamics observed in our varying concentration experiment have also been reported in other studies of doxorubicin effects on breast cancer cell lines, both as monotherapy and in combination with other therapeutic agents [52],[61],[62].

We used the parameter distributions obtained from our single-dose model fits to the varying drug concentration datasets to empirically fit various parameter formulas as functions of doxorubicin concentration, as shown in Figure 3. The model simulations generated from our proposed empirical parameter formulas were able to capture a spectrum of model solutions that encompass the dynamics observed in our data from Experiment 1 (see Figure 4). We observed clear exponentially decaying trends for the fraction of drug-resistant cells (*f*_*r*_) and the proliferation rates of drug-resistant and drug-sensitive tumor cells (*g*_*r*_ and g_*s*_, respectively) as doxorubicin concentration increases (see Figure 3a-c). These trends seem to capture the growth-inhibition effect of doxorubicin as well as the dose-response curve for this drug within our mechanistic modeling framework, in which doxorubicin efficacy has been observed to plateau at high concentrations [63],[64]. The distributions of the doxorubicin-induced death rate in drug-sensitive cells (*k*_*d*_) exhibited a non-monotonic trend as doxorubicin concentration was varied, which we approximated with a Morse-potential relationship [60]. This result was counterintuitive, as we had initially anticipated a strictly decreasing trend in *k*_*d*_ for higher doxorubicin doses, which would indicate an increasingly more intense effect of treatment-induced death. However, the cytotoxic action of doxorubicin [50]–[52] also induces cell cycle arrest. The interplay between these two drug-induced effects may ultimately lead to nonlinear tumor cell responses, such as the one captured by the empirical formula for *k*_*d*_ proposed in this work. The relative participation of cell death and cell cycle arrest in the overall doxorubicin effect on breast cancer cells may follow more complex dynamics that are not fully captured by our models, and thus requires additional investigation.

Experiment 2 involved the delivery of two doses of doxorubicin to each replicate of MCF-7 breast cancer cells at varying-inter-treatment intervals ranging from 0 to 16 days (Table 1). We fit two versions of our multiple-dose model to these datasets: either with constant parameters or with *f*_*r*_ and γ_*d*_ varied at each drug dose. The model with constant parameters sufficed to describe the observed cell dynamics for inter-treatment intervals from 0 to 6 days, while the model with varying *f*_*r*_ and γ_*d*_ was superior for inter-treatment intervals from 8 to 16 days (< 0.05, see Figure 5). For two consecutive doses of doxorubicin delivered at varying inter-treatment intervals, our results showed significantly poorer tumor control with longer inter-treatment intervals (< 0.05, see Supplementary Table S11). In the fittings from the model with varying *f*_*r*_ and γ_*d*_, we observed that the second dose induced a significantly larger *f*_*r*_ and a lower γ_*d*_ (< 0.05, see Figure 7), further supporting the adoption of an adaptive model parameterization for inter-treatment intervals from 8 to 16 days. From a biological perspective, these changes in *f*_*r*_ and γ_*d*_ suggest that longer inter-treatment intervals contribute to the development of chemoresistance in both tumor cell subpopulations in our model. Indeed, long inter-treatment intervals may allow cancer cells to acquire chemoresistance through processes like treatment-induced mutations, altered epigenetics, and phenotype switching, which ultimately limit the efficacy of the second dose and may lead to tumor regrowth [7]– [10],[17]–[19]. This phenomenon has been observed in preclinical studies [56]–[58], but the trends are less clear in the clinical setting [56],[65]–[68].

In Experiment 3, we treated MCF-7 breast cancer cells with multiple doses of doxorubicin at either 2-day or 2-week inter-treatment intervals (Table 1). We observed significantly improved tumor control with an increased number of doses delivered at a 2-day inter-treatment interval (*p* < 0.05, see Supplementary Table S15), with tumor growth effectively suppressed after two or more doxorubicin doses. When the treatment interval was extended to 2 weeks, tumor growth was significantly decelerated (*p* < 0.05, see Supplementary Table S21) but not suppressed, aligning with our previous conclusions that longer inter-treatment intervals may promote chemoresistance. Moreover, these results underscore that, in comparison to the total number of doses, it is the treatment interval that holds a critical impact on determining overall tumor control. Indeed, as most patients receive chemotherapy treatments delivered every 1-3 weeks, our results point to the clinical importance of optimizing treatment interval in designing effective drug regimens [30],[56],[65]–[68]. Additionally, the evolving distributions for the varying *f*_*r*_ and γ_*d*_ from the model fits to the 2-week inter-treatment interval datasets (see Figure 9) exhibit trends that potentially explain the relationship between the number of doses and the resulting chemoresistance dynamics. For *f*_*r*_, the initially increasing trend for the first four doses 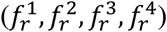, suggests a progressive increase in chemoresistance with each dose. However, the values of 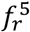 were significantly lower than those of 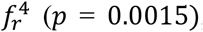, potentially indicating that increasing the number of doses may ultimately hinder chemoresistance. This is further corroborated by the trends for γ_*d*_, in which 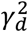 drops significantly with respect to 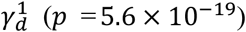, but 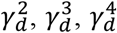, and 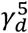 exhibit an increasing trend. This result suggests that more doses of doxorubicin can promote increasingly sensitive subpopulations that have faster transitions to treatment-induced death, thus reverting the initial chemoresistance observed in the drug-sensitive subpopulation. We do note that because we have only tested up to five doses of doxorubicin, further studies with a larger number of doses would be needed to further probe these trends.

Although our work presents promising insights into the mechanisms of chemoresistance, this study does have its limitations. First, we used a limited number of replicates within each experiment (n = 6 or 12, see Table 1). Since we do not observe uniform growth dynamics across all replicates, we would like to re-assess the observations in this study over a larger experimental setup, for example involving a higher number of replicates exposed to more diverse combinations of drug concentration, inter-treatment interval, and number of drug doses. This would enable us to investigate whether these observations are from doxorubicin effects altering tumor cell dynamics or whether the experimental conditions influence the development of a representative distribution of drug-resistant and drug-sensitive cells (e.g., ∼2,000 seeded cells/well might potentially limit the emergence of a resistant subpopulation, which may skew the observed response to treatment). Second, we also acknowledge the general limitations of extrapolating from *in vitro* systems to tumors in patients [69], as cell lines do not capture the unique, heterogeneous nature of each patient’s tumor. To address this limitation, we plan to evaluate our models on clinically-relevant breast cancer cells other than MCF-7 cells (ER+ breast cancer), such as the BT-474 (ER+HER2+ breast cancer) and MD-MBA-231 (triple-negative breast cancer) considered by Howard *et al*. [34]. Third, our cells were grown in monolayers, which are not representative of the three-dimensional tumor geometry *in vivo*. However, our mathematical models could be made readily applicable to tumor cell spheroid data. In particular, our models could be extended to a set of partial differential equations, accounting for tumor cell mobility and spatially-resolved parameters and variables, which would allow for a spatiotemporal description of spheroid growth in both *in vitro* and *in vivo* settings [30],[70]. Indeed, these extended models could incorporate other spatially-varying mechanisms beyond tumor cell dynamics, such as drug diffusion, mechanics, and angiogenesis, which have also been recognized as key components of chemoresistance and drug action [70]–[74]. Finally, we acknowledge the limitations in modeling subpopulation dynamics with total tumor cell data, and that our study thus lacks methods for specifically validating the proposed mathematical description of drug-resistant and drug-sensitive tumor cell dynamics. This issue could potentially be addressed by incorporating methods to trace cell lineage, which would enable the collection of time-resolved measurements of the diverse drug-sensitive and drug-resistant phenotypes in the tumor cell population. For instance, Al’Khafaji *et al*. have developed a functionalized lineage tracing tool to track both cell lineages and direct lineage-specific gene expression using barcoded gRNAs [75]. Then, fitting these data to an extension of our model to a multicompartment formulation describing the dynamics of the various detected drug sensitivity phenotypes could provide a more precise insight into the dose-dependent response (including refined parameter empirical formulas) and how timing and the number of doses mediate the global response of the tumor cell population.

In future studies, we intend to explore a refinement of our model to account for the mechanisms underlying the trends observed in the empirical parameter formulas from this work, which will help us further understand doxorubicin effects. Further experimentally informed studies with our mechanistic models could also contribute to identifying the optimal timing and frequency for doxorubicin delivery in preclinical scenarios. Indeed, we would like to explore optimal control theory [74],[76] *in vitro* through heterogeneous multiclonal cultures to identify optimal treatment combinations of doxorubicin concentration, treatment interval, and number of doses. Ultimately, we think that the mechanistic insights provided by our models and proposed empirical formulas could guide the identification of the minimal dose range required to effectively inhibit breast cancer growth *in vivo* and achieve optimal tumor control, which are of great clinical interest to derive minimally toxic therapeutic regimens [50],[74],[77],[78]. Thus, we believe that the complex dynamics underlying the dose-dependent effect of doxorubicin deserve further research coupling extensive experiments with mechanistic modeling.

## Conclusion

We have developed a biologically-based, mathematical model of MCF-7 breast cancer cell response to doxorubicin accounting for the development of chemoresistance, which significantly extends the experimentally-informed mechanistic models by Howard *et al*. [34]. To this end, we proposed a modeling framework that can accommodate multiple doxorubicin doses as well as an adaptive parameterization with each drug dose. We show that model fittings to longitudinal, time-resolved microscopy data of MCF-7 breast cancer cells could remarkably recapitulate the observed growth dynamics for all experimental scenarios varying in either drug concentration, inter-treatment interval, or number of doses. We also propose empirical formulas that describe model parameters as functions of doxorubicin concentration, which could contribute to refining our mechanistic model and further our understanding of doxorubicin action. We report significantly improved tumor control with higher doxorubicin concentrations, shorter inter-treatment intervals, and a higher number of doses. We also observe that longer inter-treatment intervals potentially promote chemoresistance through higher resistant fractions and delayed transitions to treatment-induced death in drug-sensitive subpopulations. Our findings show promise in furthering our understanding of doxorubicin action and chemoresistance progression, while also representing a step towards systematically exploring optimal treatment regimens *in vitro*.

## Data availability

The experimental datasets leveraged in the current study as well as the MATLAB scripts for their analysis with our mechanistic models are available at Zenodo (https://doi.org/10.5281/zenodo.5722432).

## Acknowledgments

G.L. was partially supported by a Peter O’Donnell Jr. Postdoctoral Fellowship from the Oden Institute for Computational Engineering and Sciences at The University of Texas at Austin and acknowledges funding from the European Union’s Horizon 2020 research and innovation program under the Marie Skłodowska-Curie grant agreement No. 838786. We thank the National Institutes of Health for funding through NCI R01CA240589, U01CA174706, R01 CA186193, U24 CA226110, and U01CA253540. We thank the Cancer Prevention and Research Institute of Texas for support through CPRIT RR160005; T.E.Y is a CPRIT Scholar in Cancer Research.

## Author Contributions

Conceptualization: E.Y.Y., G.L., T.E.Y.

Methodology: E.Y.Y., G.L., G.R.H., A.B., T.E.Y.

Software: E.Y.Y., G.L.

Validation: E.Y.Y., G.L.

Formal analysis: E.Y.Y., G.L., T.E.Y.

Investigation: E.Y.Y., G.L., G.R.H., A.B., T.E.Y.

Resources: G.R.H., A.B., T.E.Y.

Data curation: E.Y.Y., G.R.H., A.B.

Writing-original draft: E.Y.Y., G.L., T.E.Y.

Writing-review and editing: E.Y.Y., G.L., G.R.H., A.B., T.E.Y

Visualization: E.Y.Y., G.L.

Supervision: G.L., T.E.Y.

Project Administration: G.L.

Funding acquisition: T.E.Y.

