## Supplemental Figures and Tables for "Mathematical characterization of population dynamics in breast cancer cells treated with doxorubicin"

### Abbreviations

NRMSE: Normalized root mean square error

$R^2$: Coefficient of determination

PCC: Pearson correlation coefficient

CCC: Concordance correlation coefficient

### Supplementary Tables

| Parameter | Definition | Units |
|---|---|---|
| $\boldsymbol{N_0}$ | Initial number of tumor cells | cells |
| $\boldsymbol{g_0}$ | Baseline proliferation rate | $h^{-1}$ |
| $\boldsymbol{\theta_u}$ | Untreated carrying capacity | cells |
| $\boldsymbol{\theta_{Dox}}$ | Treated carrying capacity | cells |
| $\boldsymbol{f_r}$ | Fraction of resistant cells | - |
| $\boldsymbol{g_r}$ | Proliferation rate of drug-resistant tumor cells | $h^{-1}$ |
| $\boldsymbol{g_s}$ | Proliferation rate of drug-sensitive tumor cells | $h^{-1}$ |
| $\boldsymbol{k_d}$ | Doxorubicin-induced death rate of drug-sensitive tumor cells | $h^{-1}$ |
| $\boldsymbol{\gamma_d}$ | Doxorubicin-induced death delay rate of drug-sensitive tumor cells | $h^{-1}$ |

**Supplementary Table S1.** Definition and units of the model parameters.

| Dose [nM] | Initial guess and bounds | | | | | | | |
|---|---|---|---|---|---|---|---|---|
| | **$N_0$ [cells]** | **$g_0$ [$h^{-1}$]** | **$\theta_{Dox}$ [cells]** | **$f_r$ [ - ]** | **$g_r$ [$h^{-1}$]** | **$g_s$ [$h^{-1}$]** | **$k_d$ [$h^{-1}$]** | **$\gamma_d s$ [$h^{-1}$]** |
| **0** | $N_{0,obs}$ [700, 2000] | 0.025 [0.0075, 0.035] | 60000 [40000, 70000]* | - | - | - | - | - |
| **10** | $N_{0,obs}$ [1200, 3000] | 0.03 [0.02, 0.035] | 60000 [15000, 90000] | 0.75 [0, 1] | 0.02 [0.001, 0.05] | 0.05 [0.001, 0.065] | -0.001 [-0.01, 0] | 0.01 [1/120, 1/10] |
| **20** | 〃 | 〃 | 〃 | 〃 | 〃 | 〃 | 〃 | 〃 |
| **35** | 〃 | 〃 | 50000 [15000, 90000] | 0.5 [0, 1] | 〃 | 〃 | 〃 | 0.015 [1/120, 1/10] |
| **50** | 〃 | 〃 | 60000 [15000, 90000] | 〃 | 〃 | 〃 | 〃 | 〃 |
| **75** | 〃 | 〃 | 〃 | 〃 | 〃 | 〃 | 〃 | 0.02 [1/120, 1/10] |
| **100** | 〃 | 〃 | 〃 | 〃 | 〃 | 〃 | 〃 | 0.035 [1/120, 1/10] |
| **125** | 〃 | 〃 | 〃 | 〃 | 〃 | 0.01 [0.001, 0.065] | 〃 | 0.04 [1/120, 1/10] |
| **150** | 〃 | 〃 | 80000 [15000, 90000] | 0.25 [0, 1] | 〃 | 〃 | 〃 | 0.05 [1/120, 1/10] |
| **300** | 〃 | 〃 | 60000 [15000, 90000] | 〃 | 〃 | 0.005 [0.001, 0.065] | -0.0025 [-0.01, 0] | 〃 |

**Supplementary Table S2**. Parameter initial guesses and bounds used for fitting the single-dose model to time courses resulting from cells treated with a single dose of doxorubicin at varying concentrations (n = 6 per tested concentration; Experiment 1 in Table 1 of the main text). A ditto mark ( 〃 ) indicates that the initial guess and bounds are identical to the values in the cells above. *For 0 nM, we report the initial guess and bounds for $\theta_u$ instead of $\theta_{Dox}$.

| Dose [nM] | Parameter values | | | | | | | |
|---|---|---|---|---|---|---|---|---|
| | $N_0$ [cells] | $g_0$ [$h^{-1}$] | $\theta_{Dox}$ [cells] | $f_r$ [ - ] | $g_r$ [$h^{-1}$] | $g_s$ [$h^{-1}$] | $k_d$ [$h^{-1}$] | $\gamma_d$ [$h^{-1}$] |
| **0** | 1276<br>[1120, 1409] | 0.028<br>[0.027, 0.028] | 54640<br>[51414, 55769]* | - | - | - | - | - |
| **10** | 1400<br>[1346, 1482] | 0.026<br>[0.026, 0.027] | 63579<br>[61403, 66880] | 0.82<br>[0.80, 0.85] | 0.019<br>[0.019, 0.021] | 0.045<br>[0.04, 0.05] | $-7.6\times10^{-4}$<br>[$-9.3\times10^{-4}$, $-2.4\times10^{-4}$] | 0.01<br>[0.0084, 0.012] |
| **20** | 1444<br>[1283, 1475] | 0.026<br>[0.025, 0.027] | 77484<br>[71701, 83936] | 0.54<br>[0.50, 0.58] | 0.015<br>[0.014, 0.015] | 0.037<br>[0.031, 0.039] | -0.0054<br>[-0.0091, -0.0038] | 0.018<br>[0.015, 0.022] |
| **35** | 1346<br>[1248, 1386] | 0.024<br>[0.023, 0.024] | 63576<br>[48622, 73338] | 0.071<br>[0.049, 0.18] | 0.018<br>[0.013, 0.021] | 0.031<br>[0.029, 0.035] | -0.0072<br>[-0.008, -0.0059] | 0.02<br>[0.019, 0.027] |
| **50** | 1370<br>[1305, 1499] | 0.024<br>[0.023, 0.025] | 68827<br>[61904, 71593] | 0.051<br>[0.036, 0.091] | 0.014<br>[0.011, 0.021] | 0.027<br>[0.025, 0.035] | -0.008<br>[-0.009, -0.0067] | 0.025<br>[0.021, 0.031] |
| **75** | 1469<br>[1326, 1563] | 0.024<br>[0.022, 0.025] | 68569<br>[52398, 77716] | 0.031<br>[0.012, 0.036] | 0.011<br>[0.0081, 0.016] | 0.021<br>[0.015, 0.031] | -0.007<br>[-0.0077, -0.0061] | 0.028<br>[0.023, 0.033] |
| **100** | 1480<br>[1323, 1564] | 0.023<br>[0.022, 0.025] | 68167<br>[66466, 79285] | 0.02<br>[0.015, 0.057] | 0.0079<br>[0.0024, 0.0091] | 0.026<br>[0.011, 0.032] | -0.0055<br>[-0.0078, -0.0042] | 0.038<br>[0.025, 0.046] |
| **125** | 1435<br>[1352, 1505] | 0.026<br>[0.024, 0.028] | 68167<br>[20746, 68167] | 0.0095<br>[$9\times10^{-5}$, 0.057] | 0.0031<br>[0.001, 0.011] | 0.013<br>[0.0058, 0.018] | -0.0049<br>[-0.0086, -0.0041] | 0.024<br>[0.021, 0.046] |
| **150** | 1444<br>[1234, 1562] | 0.024<br>[0.022, 0.025] | 68167<br>[68167, 88076] | 0.0092<br>[$4\times10^{-5}$, 0.02] | 0.0035<br>[0.002, 0.015] | 0.012<br>[0.009, 0.024] | -0.0047<br>[-0.0068, -0.0043] | 0.031<br>[0.024, 0.033] |
| **300** | 1482<br>[1381, 1546] | 0.025<br>[0.024, 0.025] | 68167<br>[68167, 68167] | 0.0072<br>[$5\times10^{-4}$, 0.014] | 0.0028<br>[0.0018, 0.0063] | 0.014<br>[0.0096, 0.022] | -0.0046<br>[-0.0051, -0.0043] | 0.043<br>[0.033, 0.06] |

**Supplementary Table S3**. Median and range of the single-dose model parameters obtained from the fits to the time courses resulting from cells treated with a single dose of doxorubicin at varying concentrations (n = 6 per tested concentration; Experiment 1 in Table 1 of the main text). *For 0 nM, we report the mean and range for $\theta_u$ instead of $\theta_{Dox}$.

| Dose [nM] | Quality of fit metrics | | | |
|---|---|---|---|---|
| | **NRMSE [%]** | $\boldsymbol{R^2}$ | **PCC** | **CCC** |
| **0** | 1.11 [0.76, 1.40] | >0.999 [>0.999, >0.999] | >0.999 [>0.999, >0.999] | 0.986 [0.985, 0.986] |
| **10** | 1.13 [0.804, 1.2] | >0.999 [>0.999, >0.999] | >0.99 [>0.999, >0.999] | 0.984 [0.984, 0.985] |
| **20** | 1.46 [1.23, 2.29] | 0.999 [0.999, >0.999] | >0.99 [0.999, >0.999] | 0.986 [0.986, 0.986] |
| **35** | 2.58 [0.804, 2.87] | 0.999 [0.999, 0.999] | 0.999 [0.999, >0.999] | 0.991 [0.990, 0.991] |
| **50** | 3.07 [0.804, 4.37] | 0.998 [0.998, >0.999] | 0.999 [0.999, >0.999] | 0.991 [0.991, 0.992] |
| **75** | 2.75 [0.804, 4.61] | 0.999 [0.995, >0.999] | 0.999 [0.997, >0.999] | 0.994 [0.992, 0.994] |
| **100** | 5.19 [0.804, 10.2] | 0.993 [0.959, 0.997] | 0.996 [0.980, 0.998] | 0.991 [0.973, 0.993] |
| **125** | 4.53 [0.804, 8.48] | 0.996 [0.972, 0.997] | 0.998 [0.986, 0.999] | 0.993 [0.981, 0.994] |
| **150** | 8.48 [0.804, 12.2] | 0.990 [0.985, 0.999] | 0.995 [0.993, >0.999] | 0.991 [0.989, 0.996] |
| **300** | 9.06 [0.804, 12] | 0.990 [0.981, 0.996] | 0.995 [0.991, 0.998] | 0.992 [0.987, 0.994] |

**Supplementary Table S4**. Median and range of the quality of fit metrics for the single-dose model fits to the time courses resulting from cells treated with a single dose of doxorubicin at varying concentrations (n = 6 per tested concentration; Experiment 1 in Table 1 of the main text).

| Dose [nM] | **0** | **10** | **20** | **35** | **50** | **75** | **100** | **125** | **150** | **300** |
|---|---|---|---|---|---|---|---|---|---|---|
| **0** | - | 0.9372 | **0.0022** | **0.0087** | **0.0022** | **0.0022** | **0.0022** | **0.0022** | **0.0043** | **0.0022** |
| **10** | - | - | **0.0022** | **0.0022** | **0.0022** | **0.0022** | **0.0022** | **0.0022** | **0.0022** | **0.0022** |
| **20** | - | - | - | 0.1797 | 0.0931 | **0.0022** | **0.0022** | **0.0022** | 0.0649 | **0.0022** |
| **35** | - | - | - | - | 0.0931 | **0.0043** | **0.0022** | **0.0022** | 0.0649 | **0.0022** |
| **50** | - | - | - | - | - | 0.0649 | **0.0043** | **0.0022** | 0.0931 | **0.0022** |
| **75** | - | - | - | - | - | - | 0.1797 | **0.0152** | 0.1797 | **0.0022** |
| **100** | - | - | - | - | - | - | - | 0.0931 | 0.3939 | **0.0022** |
| **125** | - | - | - | - | - | - | - | - | 0.3939 | 0.9372 |
| **150** | - | - | - | - | - | - | - | - | - | 0.2403 |
| **300** | - | - | - | - | - | - | - | - | - | - |

**Supplementary Table S5.** P-values from two-sided Wilcoxon rank sum tests comparing the observed final tumor cell numbers for every distinct combination of the varying doxorubicin concentration datasets (n = 6 per tested concentration; Experiment 1 in Table 1 of the main text). Values bolded in red indicate $p < 0.05$.

| Parameter formula | Parameter values | | | |
|---|---|---|---|---|
| | $\alpha_1$ | $\alpha_2$ | $\alpha_3$ | $\alpha_4$ |
| $f_r(D) = \alpha_1 e^{-(\alpha_2 D)} + \alpha_3$ | 1.63<br>[1.05, 2.20] | 0.065<br>[0.038, 0.093] | 0.0032<br>[-0.061, 0.068] | - |
| $g_r(D) = \alpha_1 e^{-(\alpha_2 D)}$ | 0.022<br>[0.017, 0.026] | 0.010<br>[0.0061, 0.015] | - | - |
| $g_s(D) = \alpha_1 e^{-(\alpha_2 D)}$ | 0.043<br>[0.033, 0.053] | 0.0074<br>[0.0036, 0.011] | - | - |
| $k_d(D) = \alpha_1 (1 - e^{-(\alpha_2 (D - \alpha_3))})^2 - \alpha_4$ | 0.0033<br>[0.0023, 0.0044] | 0.028<br>[0.02, 0.035] | 41.92<br>[35.24, 48.60] | 0.0078<br>[0.0072, 0.0084] |
| $\gamma_d(D) = \alpha_1 - \alpha_2 e^{-(\alpha_3 D)}$ | 0.035<br>[0.027, 0.043] | 0.026<br>[0.006, 0.047] | 0.0104<br>[-0.0062, 0.0270] | - |

**Supplementary Table S6**. Fitted empirical parameter values and the corresponding 95% confidence intervals for the empirical parameter formulas derived from the single-dose model fits to the varying doxorubicin concentration datasets (n = 6 per tested concentration; Experiment 1 in Table 1 of the main text).

| Parameter formula | Quality of fit metrics | | | |
|---|---|---|---|---|
| | **NRMSE [%]** | $R^2$ | **PCC** | **CCC** |
| $f_r(D) = \alpha_1 e^{-(\alpha_2 D)} + \alpha_3$ | 153.64* | 0.977 | 0.985 | 0.876 |
| $g_r(D) = \alpha_1 e^{-(\alpha_2 D)}$ | 16.73 | 0.909 | 0.953 | 0.845 |
| $g_s(D) = \alpha_1 e^{-(\alpha_2 D)}$ | 17.65 | 0.818 | 0.912 | 0.810 |
| $k_d(D) = \alpha_1 (1 - e^{-(\alpha_2 (D - \alpha_3))})^2 - \alpha_4$ | 5.45 | 0.978 | 0.992 | 0.879 |
| $\gamma_d(D) = \alpha_1 - \alpha_2 e^{-(\alpha_3 D)}$ | 17.94 | 0.784 | 0.883 | 0.779 |

**Supplementary Table S7**. Quality of fit metrics for the empirical parameter formulas derived from the single-dose model fits to the varying doxorubicin concentration datasets (n = 6 per tested concentration; Experiment 1 in Table 1 of the main text). *For larger drug concentrations, $f_r$ takes on values on the order of $10^{-5}$, such that natural variations of $f_r$ across replicates may result in NRMSE values greater than 100%. However, these variations have a negligible impact on model outcome.

| Interval [d] | Initial guess and bounds | | | | | | | | | |
|---|---|---|---|---|---|---|---|---|---|---|
| | **$N_0$ [cells]** | **$g_0$ [$h^{-1}$]** | **$\theta_{Dox}$ [cells]** | **$f_r^1$ [ - ]** | **$f_r^2$ [ - ]** | **$g_r$ [$h^{-1}$]** | **$g_s$ [$h^{-1}$]** | **$k_d$ [$h^{-1}$]** | **$\gamma_d^1$ [$h^{-1}$]** | **$\gamma_d^2$ [$h^{-1}$]** |
| **Multiple-dose model with constant parameters** | | | | | | | | | | |
| **0** | $N_{0,obs}$ [1000, 2500] | 0.0234 [0.02, 0.035] | 60000 [30000, 90000] | 0.0337 [0, 1] | | 0.0101 [0.001, 0.05] | 0.0277 [0.001, 0.05] | -0.0069 [-0.01, 0] | 0.0325 [0, 1/15] | |
| **2** | 〃 | 〃 | 〃 | 〃 | | 〃 | 〃 | 〃 | 〃 | |
| **4** | 〃 | 〃 | 〃 | 〃 | | 〃 | 〃 | 〃 | 〃 | |
| **6** | 〃 | 〃 | 〃 | 〃 | | 〃 | 〃 | 〃 | 〃 | |
| **Multiple-dose model with varying $f_r$ and $\gamma_d$** | | | | | | | | | | |
| **8** | $N_{0,obs}$ [1200, 2500] | 0.0234 [0.02, 0.035] | 60000 [15000, 90000] | 0.0337 [0, 0.1] | 0.005 [0, 1] | 0.0101 [0.001, 0.05] | 0.0277 [0.001, 0.055] | $-5\times10^{-4}$ [-0.01, 0] | 0.0325 [0, 1/15] | 1/75 [0, 1/15] |
| **10** | 〃 | 〃 | 〃 | 〃 | 〃 | 〃 | 〃 | 〃 | 〃 | 〃 |
| **12** | 〃 | 〃 | 〃 | 〃 | 〃 | 〃 | 〃 | 〃 | 〃 | 〃 |
| **14** | 〃 | 〃 | 〃 | 〃 | 〃 | 〃 | 〃 | 〃 | 〃 | 〃 |
| **16** | 〃 | 〃 | 〃 | 〃 | 〃 | 〃 | 〃 | 〃 | 〃 | 〃 |

**Supplementary Table S8**. Parameter initial guesses and bounds used for fitting the multiple-dose model to the time courses resulting from cells treated with two consecutive doses of 75 nM doxorubicin delivered at varying inter-treatment intervals (n = 12 per tested interval; Experiment 2 in Table 1 of the main text). A ditto mark ( 〃 ) indicates that the initial guess and bounds are identical to the values in the cells above.

| Interval [d] | Parameter values | | | | | | | | | |
|---|---|---|---|---|---|---|---|---|---|---|
| | $N_0$ [cells] | $g_0$ [$h^{-1}$] | $\theta_{Dox}$ [cells] | $f_r^1$ [ - ] | $f_r^2$ [ - ] | $g_r$ [$h^{-1}$] | $g_s$ [$h^{-1}$] | $k_d$ [$h^{-1}$] | $\gamma_d^1$ [$h^{-1}$] | $\gamma_d^2$ [$h^{-1}$] |
| **Multiple-dose model with constant parameters** | | | | | | | | | | |
| **0** | 1468<br>[1354, 1542] | 0.024<br>[0.023, 0.025] | 53376<br>[42605, 80016] | 0.0019<br>[$7\times10^{-5}$, 0.017] | | 0.002<br>[0.001, 0.014] | 0.022<br>[0.012, 0.026] | -0.0035<br>[-0.0047, -0.002] | 0.046<br>[0.027, 0.057] | |
| **2** | 1468<br>[1381, 1591] | 0.024<br>[0.024, 0.025] | 53376<br>[53376, 53376] | 0.012<br>[0.0069, 0.23] | | 0.0045<br>[0.0028, 0.0069] | 0.026<br>[0.023, 0.038] | -0.0035<br>[-0.0047, -0.0032] | 0.048<br>[0.042, 0.063] | |
| **4** | 1514<br>[1396, 1572] | 0.024<br>[0.023, 0.025] | 53376<br>[53376, 53376] | 0.018<br>[0.016, 0.084] | | 0.0067<br>[0.0054, 0.0079] | 0.022<br>[0.021, 0.025] | -0.0037<br>[-0.0042, -0.0034] | 0.04<br>[0.035, 0.05] | |
| **6** | 1411<br>[1312, 1591] | 0.024<br>[0.023, 0.025] | 53376<br>[30690, 53376] | 0.031<br>[0.011, 0.09] | | 0.0086<br>[0.0063, 0.011] | 0.019<br>[0.017, 0.022] | -0.0042<br>[-0.0061, -0.0033] | 0.031<br>[0.027, 0.04] | |
| **Multiple-dose model with varying $f_r$ and $\gamma_d$** | | | | | | | | | | |
| **8** | 1572<br>[1289, 1676] | 0.021<br>[0.02, 0.024] | 53376<br>[36130, 56316] | 0.0170<br>[0.0009, 0.0310] | 0.1100<br>[0.029, 0.20] | 0.0093<br>[0.0010, 0.0160] | 0.0020<br>[0.0160, 0.0230] | -0.0036<br>[-0.0043, -0.0027] | 0.0340<br>[0.0280, 0.0440] | 0.0300<br>[0.0120, 0.0440] |
| **10** | 1573<br>[1382, 1687] | 0.022<br>[0.02, 0.025] | 53376<br>[36130, 56316] | 0.021<br>[0.0055, 0.037] | 0.13<br>[0.051, 0.23] | 0.0094<br>[0.0077, 0.015] | 0.019<br>[0.015, 0.022] | -0.0034<br>[-0.0046, -0.0027] | 0.032<br>[0.024, 0.039] | 0.024<br>[0.014, 0.034] |
| **12** | 1372<br>[1209, 1479] | 0.022<br>[0.020, 0.025] | 53376<br>[46045, 67004] | 0.016<br>[0.0083, 0.033] | 0.16<br>[0.079, 0.24] | 0.0098<br>[0.0084, 0.012] | 0.02<br>[0.017, 0.024] | -0.0031<br>[-0.0036, -0.0021] | 0.041<br>[0.035, 0.051] | 0.016<br>[0.0089, 0.032] |
| **14** | 1428<br>[1327, 1850] | 0.022<br>[0.020, 0.025] | 58091<br>[37433, 73197] | 0.025<br>[0.0066, 0.042] | 0.082<br>[0.046, 0.26] | 0.01<br>[0.0078, 0.012] | 0.019<br>[0.0038, 0.024] | -0.0031<br>[-0.0048, -0.0017] | 0.042<br>[0.025, 0.065] | 0.014<br>[0.011, 0.026] |
| **16** | 1560<br>[1350, 1684] | 0.023<br>[0.021, 0.025] | 53376<br>[42964, 81236] | 0.032<br>[0.016, 0.04] | 0.13<br>[0.075, 0.25] | 0.01<br>[0.007, 0.012] | 0.019<br>[0.013, 0.021] | -0.0039<br>[-0.0049, -0.0031] | 0.04<br>[0.028, 0.047] | 0.018<br>[0.012, 0.038] |

**Supplementary Table S9**. Median and range of the multiple-dose model parameters obtained from the fits to the time courses resulting from cells treated with two consecutive doses of 75 nM doxorubicin delivered at varying inter-treatment intervals (n = 12 per tested interval; Experiment 2 in Table 1 of the main text).

| Interval [d] | Quality of fit metrics | | | |
|---|---|---|---|---|
| | **NRMSE [%]** | $R^2$ | **PCC** | **CCC** |
| **Multiple-dose model with constant parameters** | | | | |
| **0** | 4.9 [3.2, 13.5] | 0.995 [0.948, 0.998] | 0.998 [0.974, 0.999] | 0.994 [0.969, 0.995] |
| **2** | 4.39 [3.26, 4.76] | 0.996 [0.992, 0.998] | 0.998 [0.996, 0.999] | 0.994 [0.992, 0.995] |
| **4** | 6.12 [5.03, 7.27] | 0.991 [0.979, 0.994] | 0.995 [0.990, 0.997] | 0.991 [0.986, 0.993] |
| **6** | 5.27 [2.74, 9.47] | 0.992 [0.971, 0.999] | 0.996 [0.986, >0.999] | 0.992 [0.982, 0.996] |
| **Multiple-dose model with varying $f_r$ and $\gamma_d$** | | | | |
| **8** | 4.57 [3.59, 6.32] | 0.993 [0.983, 0.997] | 0.997 [0.992, 0.999] | 0.992 [0.988, 0.995] |
| **10** | 4.08 [3.59, 6.57] | 0.992 [0.974, 0.998] | 0.996 [0.988, 0.999] | 0.992 [0.983, 0.995] |
| **12** | 4.54 [3.44, 10.2] | 0.995 [0.963, 0.998] | 0.998 [0.982, 0.999] | 0.994 [0.977, 0.995] |
| **14** | 4.39 [2.79, 14.3] | 0.996 [0.769, 0.999] | 0.998 [0.911, >0.999] | 0.994 [0.898, 0.996] |
| **16** | 4.38 [3.12, 7.04] | 0.996 [0.971, 0.998] | 0.998 [0.986, 0.999] | 0.994 [0.981, 0.995] |

**Supplementary Table S10**. Median and range of the quality of fit metrics for the multiple-dose model fits to the time courses resulting from cells treated with two consecutive doses of 75 nM doxorubicin delivered at varying inter-treatment intervals (n = 12 per tested interval; Experiment 2 in Table 1 of the main text).

| Interval [d] | **0** | **2** | **4** | **6** | **8** | **10** | **12** | **14** | **16** |
|---|---|---|---|---|---|---|---|---|---|
| **0** | - | 0.2145 | 0.4705 | 0.0999 | 0.0783 | **0.0226** | **0.0166** | **0.0102** | **0.0061** |
| **2** | - | - | **0.0029** | **$1.96\times10^{-4}$** | **$1.55\times10^{-4}$** | **$3.66\times10^{-5}$** | **$3.66\times10^{-5}$** | **$3.66\times10^{-5}$** | **$3.66\times10^{-5}$** |
| **4** | - | - | - | **0.0102** | **0.0029** | **$3.66\times10^{-5}$** | **$3.66\times10^{-5}$** | **$3.66\times10^{-5}$** | **$3.66\times10^{-5}$** |
| **6** | - | - | - | - | 0.5067 | **0.0262** | **0.0035** | **0.0120** | **0.0011** |
| **8** | - | - | - | - | - | 0.1749 | 0.0783 | **0.0404** | **0.0024** |
| **10** | - | - | - | - | - | - | 0.5834 | 0.4705 | 0.0690 |
| **12** | - | - | - | - | - | - | - | 0.5444 | 0.1124 |
| **14** | - | - | - | - | - | - | - | - | 0.5444 |
| **16** | - | - | - | - | - | - | - | - | - |

**Supplementary Table S11.** P-values from two-sided Wilcoxon rank sum tests comparing the observed final tumor cell numbers for every distinct combination of the varying inter-treatment interval datasets (n = 12 per tested interval; Experiment 2 in Table 1 of the main text). Values bolded in red indicate $p < 0.05$.

| Number of doses | Initial guess and bounds: 2-day inter-treatment interval | | | | | | | |
|---|---|---|---|---|---|---|---|---|
| | $N_0$ [cells] | $g_0$ [$h^{-1}$] | $\theta_{Dox}$ [cells] | $f_r$ [ - ] | $g_r$ [$h^{-1}$] | $g_s$ [$h^{-1}$] | $k_d$ [$h^{-1}$] | $\gamma_d$ [$h^{-1}$] |
| **1** | $N_{0,obs}$<br>[1000, 2500] | 0.0234<br>[0.0225, 0.05] | 60000<br>[20000, 90000] | 0.0337<br>[0, 0.05] | 0.0101<br>[$1\times10^{-4}$, 0.05] | 0.0277<br>[$1\times10^{-4}$, 0.05] | $-1\times10^{-5}$<br>[-0.05, 0] | 0.0325<br>[0, 1] |
| **2** | 〃 | 〃 | 〃 | 0.01<br>[0, 0.05] | 0.0025<br>[$1\times10^{-4}$, 0.05] | 0.0025<br>[$1\times10^{-4}$, 0.05] | -0.015<br>[-0.05, 0] | 〃 |
| **3** | 〃 | 〃 | 〃 | 0.001<br>[0, 0.05] | 〃 | 〃 | 〃 | 0.05<br>[0, 1] |
| **4** | 〃 | 〃 | 〃 | $1\times10^{-5}$<br>[0, 0.05] | 〃 | 〃 | 〃 | 0.08<br>[0, 1] |
| **5** | 〃 | 〃 | 〃 | $1\times10^{-9}$<br>[0, 0.05] | $1.5\times10^{-4}$<br>[$1\times10^{-4}$, 0.05] | $1.5\times10^{-4}$<br>[$1\times10^{-4}$, 0.05] | -0.035<br>[-0.05, 0] | 0.75<br>[0, 1] |

**Supplementary Table S12**. Parameter initial guesses and bounds used for fitting the multiple-dose model to the time courses resulting from cells treated with a varying number of 75 nM doxorubicin doses delivered at 2-day inter-treatment intervals (n = 12 per tested number of doses; Experiment 3 in Table 1 of the main text). A ditto mark ( 〃 ) indicates that the initial guess and bounds are identical to the values in the cells above.

| Number of Doses | Parameter values: 2-day inter-treatment interval | | | | | | | |
|---|---|---|---|---|---|---|---|---|
| | $N_0$ [cells] | $g_0$ [$h^{-1}$] | $\theta_{Dox}$ [cells] | $f_r$ [ - ] | $g_r$ [$h^{-1}$] | $g_s$ [$h^{-1}$] | $k_d$ [$h^{-1}$] | $\gamma_d$ [$h^{-1}$] |
| **1** | 1477<br>[1276, 1712] | 0.024<br>[0.023, 0.028] | 59448<br>[34132, 87537] | 0.02<br>[0.004, 0.029] | 0.01<br>[0.0084, 0.015] | 0.021<br>[0.0088, 0.03] | -0.006<br>[-0.0089, -0.003] | 0.037<br>[0.014, 0.064] |
| **2** | 1313<br>[1194, 1503] | 0.028<br>[0.027, 0.029] | 59448<br>[59448, 59448] | 0.0088<br>[$1\times10^{-7}$, 0.05] | 0.0062<br>[0.0047, 0.0079] | 0.019<br>[0.012, 0.039] | -0.004<br>[-0.0042, -0.0034] | 0.08<br>[0.059, 0.2] |
| **3** | 1278<br>[1081, 1456] | 0.028<br>[0.026, 0.029] | 59448<br>[59448, 59448] | 0.0066<br>[$1\times10^{-7}$, 0.016] | 0.013<br>[0.0099, 0.016] | 0.029<br>[0.017, 0.038] | -0.005<br>[-0.005, -0.0041] | 0.13<br>[0.088, 0.15] |
| **4** | 1196<br>[1032, 1380] | 0.028<br>[0.026, 0.029] | 59448<br>[59448, 59448] | $6\times10^{-5}$<br>[$2\times10^{-14}$, 0.0086] | $1\times10^{-4}$<br>[$1\times10^{-4}$, 0.017] | 0.027<br>[0.023, 0.033] | -0.005<br>[-0.0058, -0.0045] | 0.11<br>[0.074, 0.15] |
| **5** | 1140<br>[1051, 1305] | 0.028<br>[0.026, 0.029] | 59448<br>[59448, 59448] | 0.031<br>[0.023, 0.047] | 0.015<br>[0.013, 0.017] | 0.035<br>[0.028, 0.05] | -0.006<br>[-0.0068, -0.0061] | 0.11<br>[0.086, 0.18] |

**Supplementary Table S13**. Median and range of the multiple-dose model parameters obtained from the fits to the time courses resulting from cells treated with a varying number of 75 nM doxorubicin doses delivered at 2-day inter-treatment intervals (n = 12 per tested number of doses; Experiment 3 in Table 1 of the main text).

| Number of doses | Quality of fit metrics: 2-day inter-treatment interval | | | |
|---|---|---|---|---|
| | **NRMSE [%]** | ***$R^2$*** | **PCC** | **CCC** |
| **1** | 3.5 [2.72, 5.79] | 0.998 [0.992, 0.999] | 0.999 [0.996, >0.999] | 0.993 [0.99, 0.994] |
| **2** | 4.72 [4.14, 19.1] | 0.997 [0.992, 0.999] | 0.998 [0.934, 0.999] | 0.994 [0.928, 0.995] |
| **3** | 12.1 [10.1, 13.7] | 0.985 [0.992, 0.999] | 0.993 [0.991, 0.995] | 0.988 [0.987, 0.991] |
| **4** | 16.2 [15.1, 17.5] | 0.975 [0.992, 0.999] | 0.989 [0.987, 0.991] | 0.984 [0.981, 0.986] |
| **5** | 14.8 [13, 16.4] | 0.983 [0.992, 0.999] | 0.993 [0.991, 0.995] | 0.988 [0.986, 0.99] |

**Supplementary Table S14.** Median and range of the quality of fit metrics for the multiple-dose model fits to the time courses resulting from cells treated with a varying number of 75 nM doxorubicin doses delivered at 2-day inter-treatment intervals (n = 12 per tested number of doses; Experiment 3 in Table 1 of the main text).

| Number of doses | **1** | **2** | **3** | **4** | **5** |
|---|---|---|---|---|---|
| **1** | **-** | **$3.66\times10^{-5}$** | **$3.64\times10^{-5}$** | **$3.64\times10^{-5}$** | **$3.64\times10^{-5}$** |
| **2** | **-** | **-** | **0.0165** | **0.0110** | **$4.68\times10^{-5}$** |
| **3** | **-** | **-** | **-** | 0.7289 | **$1.08\times10^{-4}$** |
| **4** | **-** | **-** | **-** | **-** | **0.0079** |
| **5** | **-** | **-** | **-** | **-** | **-** |

**Supplementary Table S15.** P-values from 2-sided Wilcoxon rank sum tests comparing the final tumor cell numbers for every distinct combination of the varying dose number datasets with a 2-day inter-treatment interval. (n = 12 per tested number of doses; Experiment 3 in Table 1 of the main text). Values bolded in red indicate $p < 0.05$.

| Number of Doses | Initial guess and bounds: 2-week inter-treatment interval | | | | | |
|---|---|---|---|---|---|---|
| | $\boldsymbol{N_0}$ **[cells]** | $\boldsymbol{g_0}$ **[h$^{-1}$]** | $\boldsymbol{\theta_{Dox}}$ **[cells]** | $\boldsymbol{g_r}$ **[h$^{-1}$]** | $\boldsymbol{g_s}$ **[h$^{-1}$]** | $\boldsymbol{k_d}$ **[h$^{-1}$]** |
| **1** | $N_{0,obs}$<br>[900, 3000] | 0.0275<br>[0.02, 0.05] | 80000<br>[15000, 90000] | 0.02<br>[0.001, 0.05] | 0.0275<br>[0.0125, 0.075] | -0.0075<br>[-0.01, -0.001] |
| **2** | 〃 | 〃 | 〃 | 〃 | 〃 | 〃 |
| **3** | 〃 | 〃 | 〃 | 〃 | 〃 | 〃 |
| **4** | 〃 | 〃 | 〃 | 〃 | 〃 | 〃 |
| **5** | 〃 | 〃 | 〃 | 〃 | 〃 | 〃 |

**Supplementary Table S16**. Initial guesses and bounds of the constant parameters used for fitting the multiple-dose model to the time courses resulting from cells treated with a varying number of 75 nM doxorubicin doses delivered at 2-week inter-treatment intervals (n = 12 per tested number of doses; Experiment 3 in Table 1 of the main text). A ditto mark ( 〃 ) indicates that the initial guess and bounds are identical to the values in the cells above.

| Initial guess & bounds: 2-week inter-treatment interval | | | | |
|---|---|---|---|---|
| $\boldsymbol{f_r^1}$ **[ - ]** | $\boldsymbol{f_r^2}$ **[ - ]** | $\boldsymbol{f_r^3}$ **[ - ]** | $\boldsymbol{f_r^4}$ **[ - ]** | $\boldsymbol{f_r^5}$ **[ - ]** |
| 0.001<br>[0, 1] | 0.005<br>[0, 0.5] | 0.005<br>[0, 0.5] | 0.005<br>[0, 0.5] | 0.005<br>[0, 0.5] |
| $\boldsymbol{\gamma_d^1}$ **[h$^{-1}$]** | $\boldsymbol{\gamma_d^2}$ **[h$^{-1}$]** | $\boldsymbol{\gamma_d^3}$ **[h$^{-1}$]** | $\boldsymbol{\gamma_d^4}$ **[h$^{-1}$]** | $\boldsymbol{\gamma_d^5}$ **[h$^{-1}$]** |
| 1/50<br>[0, 1/15] | 1/50<br>[0, 1/15] | 1/50<br>[0, 1/15] | 1/50<br>[0, 1/15] | 1/50<br>[0, 1/15] |

**Supplementary Table S17**. Initial guesses and bounds of $f_r$ and $\gamma_d$ for each doxorubicin dose used for fitting the multiple-dose model to the time courses resulting from cells treated with a varying number of 75 nM doxorubicin doses delivered at 2-week inter-treatment intervals (n = 12 per tested number of doses; Experiment 3 in Table 1 of the main text). A ditto mark ( 〃 ) indicates that the initial guess and bounds are identical to the values in the cells above.

| Number of doses | Parameter values: 2-week inter-treatment interval | | | | | |
|---|---|---|---|---|---|---|
| | $N_0$ [cells] | $g_0$ [$h^{-1}$] | $\theta_{Dox}$ [cells] | $g_r$ [$h^{-1}$] | $g_s$ [$h^{-1}$] | $k_d$ [$h^{-1}$] |
| **1** | 1626<br>[1437, 1791] | 0.024<br>[0.023, 0.025] | 52460<br>[41191, 73604] | 0.013<br>[0.011, 0.015] | 0.02<br>[0.014, 0.023] | -0.0053<br>[-0.0073, -0.0032] |
| **2** | 1322<br>[1066, 1568] | 0.026<br>[0.024, 0.027] | 55086<br>[32671, 62376] | 0.015<br>[0.013, 0.018] | 0.014<br>[0.013, 0.016] | -0.0026<br>[-0.0037, -0.0023] |
| **3** | 1345<br>[1051, 1553] | 0.025<br>[0.022, 0.027] | 68390<br>[62376, 78206] | 0.018<br>[0.016, 0.019] | 0.014<br>[0.013, 0.023] | -0.0023<br>[-0.0029, -0.001] |
| **4** | 1182<br>[1017, 1417] | 0.024<br>[0.021, 0.026] | 71551<br>[54867, 77778] | 0.018<br>[0.016, 0.02] | 0.013<br>[0.013, 0.018] | -0.0014<br>[-0.0021, -0.001] |
| **5** | 1114<br>[901, 1287] | 0.024<br>[0.021, 0.025] | 70227<br>[57909, 78940] | 0.019<br>[0.016, 0.019] | 0.015<br>[0.013, 0.019] | -0.0014<br>[-0.003, -0.001] |

**Supplementary Table S18**. Median and range of the constant model parameters from the multiple-dose model fits to the time courses resulting from cells treated with a varying number of 75 nM doxorubicin doses delivered at 2-week inter-treatment intervals (n = 12 per tested number of doses; Experiment 3 in Table 1 of the main text).

| Parameter values: 2-week inter-treatment interval | | | | |
|---|---|---|---|---|
| $f_r^1$ [ - ] | $f_r^2$ [ - ] | $f_r^3$ [ - ] | $f_r^4$ [ - ] | $f_r^5$ [ - ] |
| 0.003<br>[$2\times10^{-4}$, 0.057] | 0.0076<br>[0.0032, 0.07] | 0.0076<br>[0.0032, 0.07] | 0.0076<br>[0.0032, 0.07] | 0.0076<br>[0.0032, 0.07] |
| $\gamma_d^1$ [$h^{-1}$] | $\gamma_d^2$ [$h^{-1}$] | $\gamma_d^3$ [$h^{-1}$] | $\gamma_d^4$ [$h^{-1}$] | $\gamma_d^5$ [$h^{-1}$] |
| 0.039<br>[0.02, 0.067] | 0.0097<br>[0.0049, 0.017] | 0.0097<br>[0.0049, 0.017] | 0.0097<br>[0.0049, 0.017] | 0.0097<br>[0.0049, 0.017] |

**Supplementary Table S19**. Median and range of $f_r$ and $\gamma_d$ for each doxorubicin dose from the multiple-dose model fits to the time courses resulting from cells treated with a varying number of 75 nM doxorubicin doses delivered at 2-week inter-treatment intervals (n = 12 per tested number of doses; Experiment 3 in Table 1 of the main text).

| Number of doses | Quality of fit metrics: 2-week inter-treatment interval | | | |
|---|---|---|---|---|
| | **NRMSE [%]** | $R^2$ | **PCC** | **CCC** |
| **1** | 2.38 [2.05, 3.11] | 0.999 [0.998, >0.999] | >0.999 [0.999, >0.999] | 0.993 [0.993, 0.994] |
| **2** | 3.45 [2.13, 8.57] | 0.996 [0.932, 0.999] | 0.998 [0.969, 0.999] | 0.993 [0.959, 0.995] |
| **3** | 3.69 [2.90, 6.80] | 0.993 [0.952, 0.997] | 0.997 [0.978, 0.999] | 0.992 [0.971, 0.995] |
| **4** | 3.51 [1.91, 7.02] | 0.996 [0.975, 0.999] | 0.998 [0.988, >0.999] | 0.995 [0.984, 0.997] |
| **5** | 3.10 [1.99, 6.40] | 0.996 [0.934, 0.999] | 0.998 [0.969, 0.999] | 0.995 [0.963, 0.997] |

**Supplementary Table S20.** Median and range of the quality of fit metrics for the multiple-dose model fits to time courses resulting from cells treated with a varying number of 75 nM doxorubicin doses delivered at 2-week inter-treatment intervals (n = 12 per tested number of doses; Experiment 3 in Table 1 of the main text).

| Number of doses | **1** | **2** | **3** | **4** | **5** |
|---|---|---|---|---|---|
| **1** | - | **$9.73\times10^{-5}$** | **$4.69\times10^{-5}$** | **$2.46\times10^{-5}$** | **$7.66\times10^{-5}$** |
| **2** | - | - | 0.4357 | 0.7508 | 0.5444 |
| **3** | - | - | - | 0.1572 | 0.4025 |
| **4** | - | - | - | - | 0.3708 |
| **5** | - | - | - | - | - |

**Supplementary Table S21.** P-values from 2-sided Wilcoxon rank sum tests comparing the final tumor cell numbers for every distinct combination of the varying dose number datasets with a 2-week treatment interval. (n = 12 per tested number of doses; Experiment 3 in Table 1 of the main text). Values bolded in red indicate $p < 0.05$.

#### Supplementary Figures

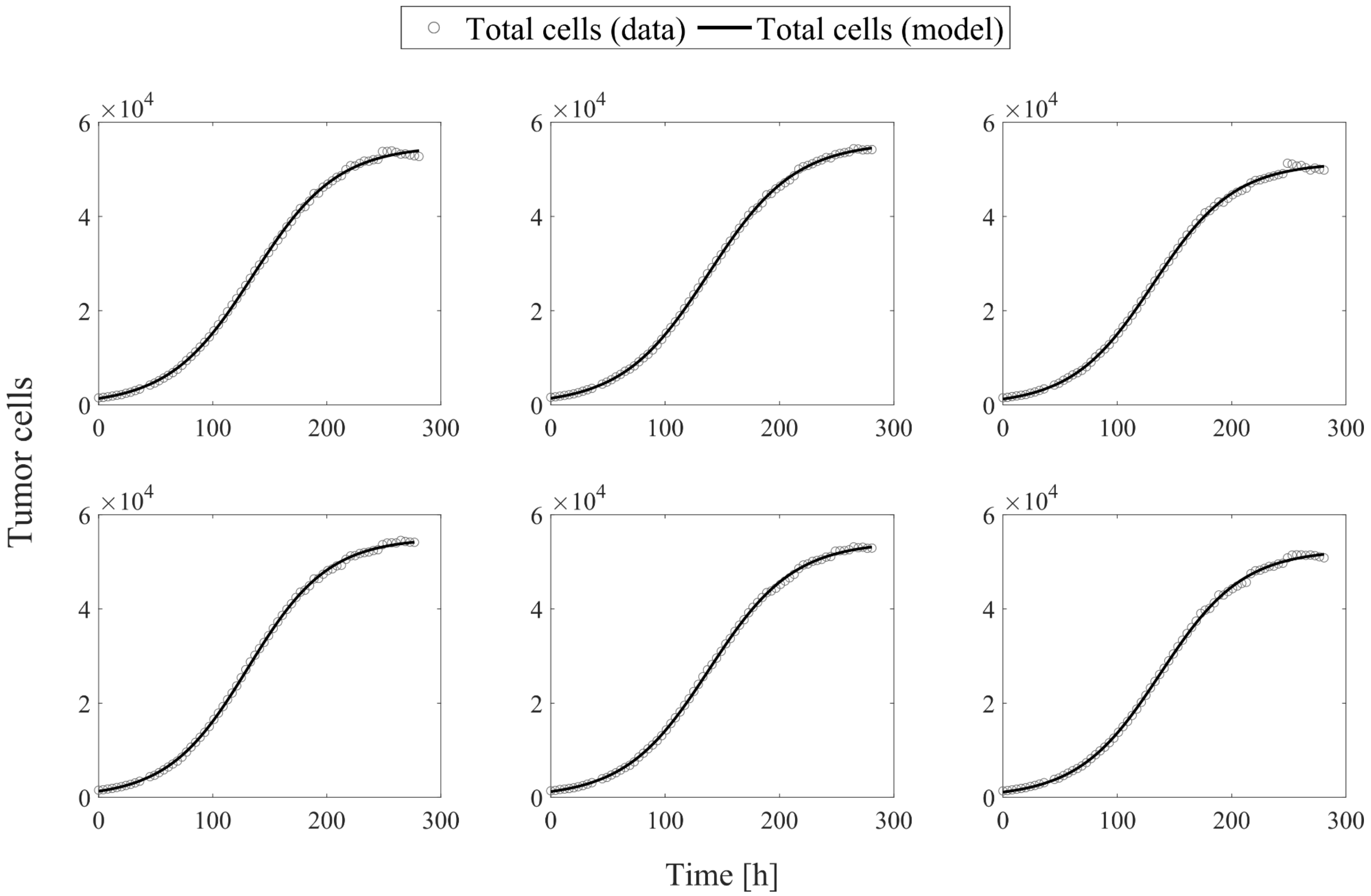

**Supplementary Figure S1.** Logistic growth model fits for all of the untreated tumor cell time courses (i.e., for a doxorubicin concentration of 0 nM in Experiment 1 in Table 1 of the main text; n = 6).

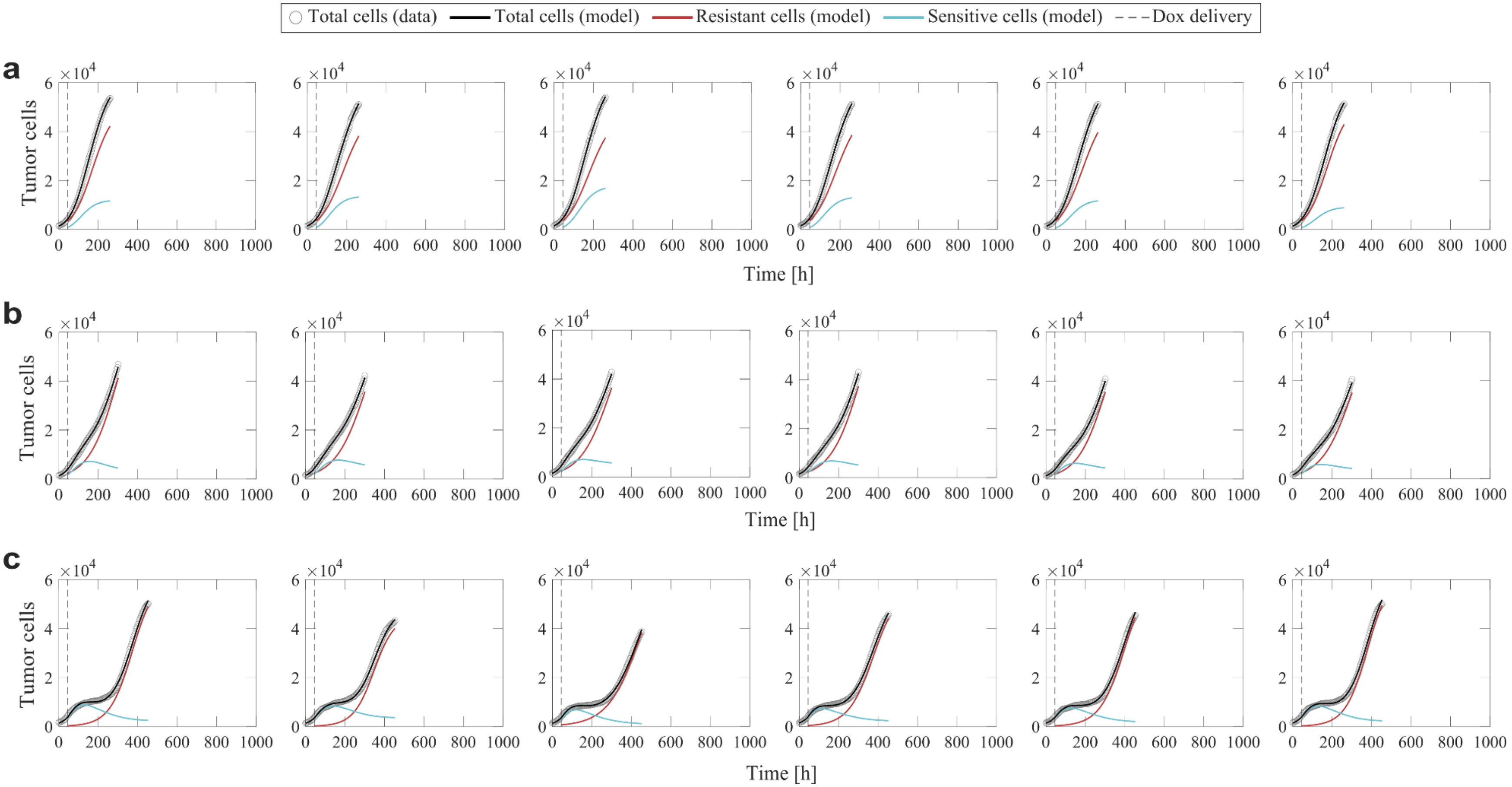

**Supplementary Figure S2.** Single-dose model fits for all of the time courses resulting from cells treated with a single dose of doxorubicin at concentrations ranging from 10 to 300 nM (n = 6 for each concentration; Experiment 1 in Table 1 of the main text). (**a**) 10 nM; (**b**) 20 nM; (**c**) 35 nM.

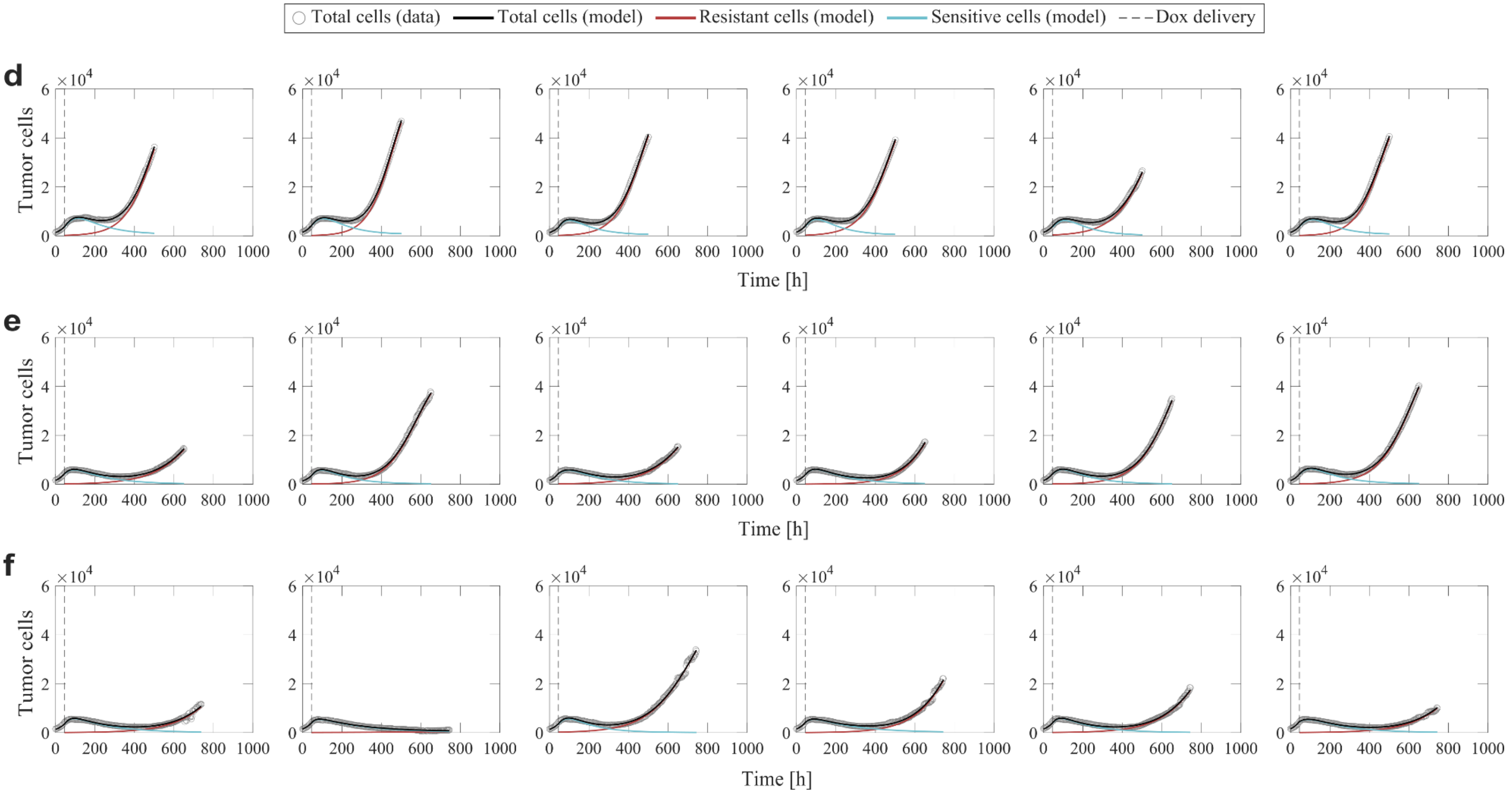

**Supplementary Figure S2 (continued).** Single-dose model fits for all of the time courses resulting from cells treated with a single dose of doxorubicin at concentrations ranging from 10 to 300 nM (n = 6 for each concentration; Experiment 1 in Table 1 of the main text). (**d**) 50nM; (**e**) 75nM; (**f**) 100 nM.

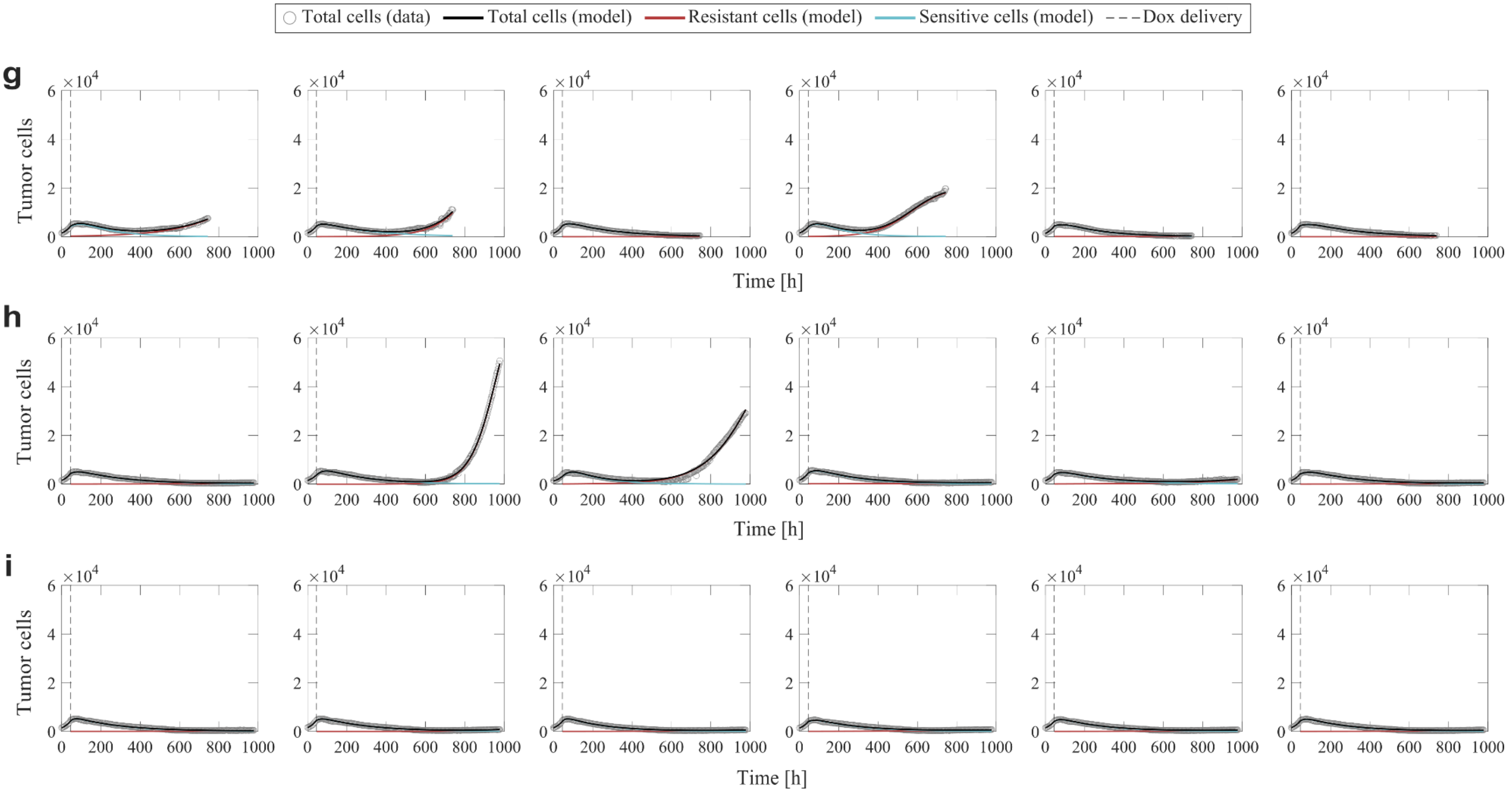

**Supplementary Figure S2 (continued).** Single-dose model fits for all of the time courses resulting from cells treated with a single dose of doxorubicin at concentrations ranging from 10 to 300 nM (n = 6 for each concentration; Experiment 1 in Table 1 of the main text). (**g**) 125 nM; (**h**) 150 nM; (**i**) 300 nM.

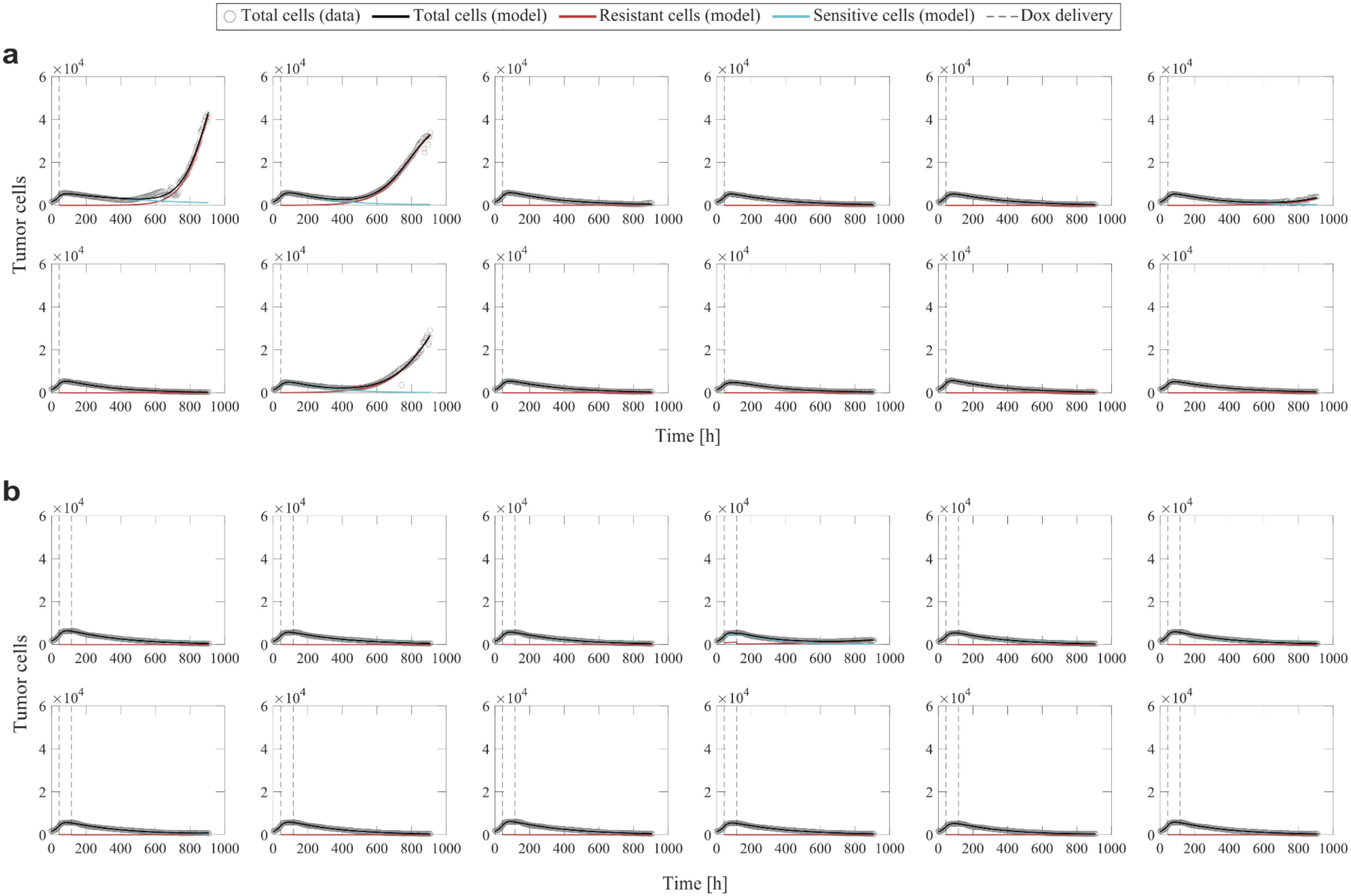

**Supplementary Figure S3.** Multiple-dose model fits for all of the time courses resulting from cells treated with two consecutive doses of 75 nM doxorubicin delivered at varying inter-treatment intervals (n = 12 for each interval; Experiment 2 in Table 1 of the main text). (**a**) 0 days; (**b**) 2 days. For (**a**)-(**d**), the multiple-dose model with constant parameters was used to fit the datasets, while for (**e**)-(**i**), the multiple-dose model with varying $f_r$ and $\gamma_d$ was used.

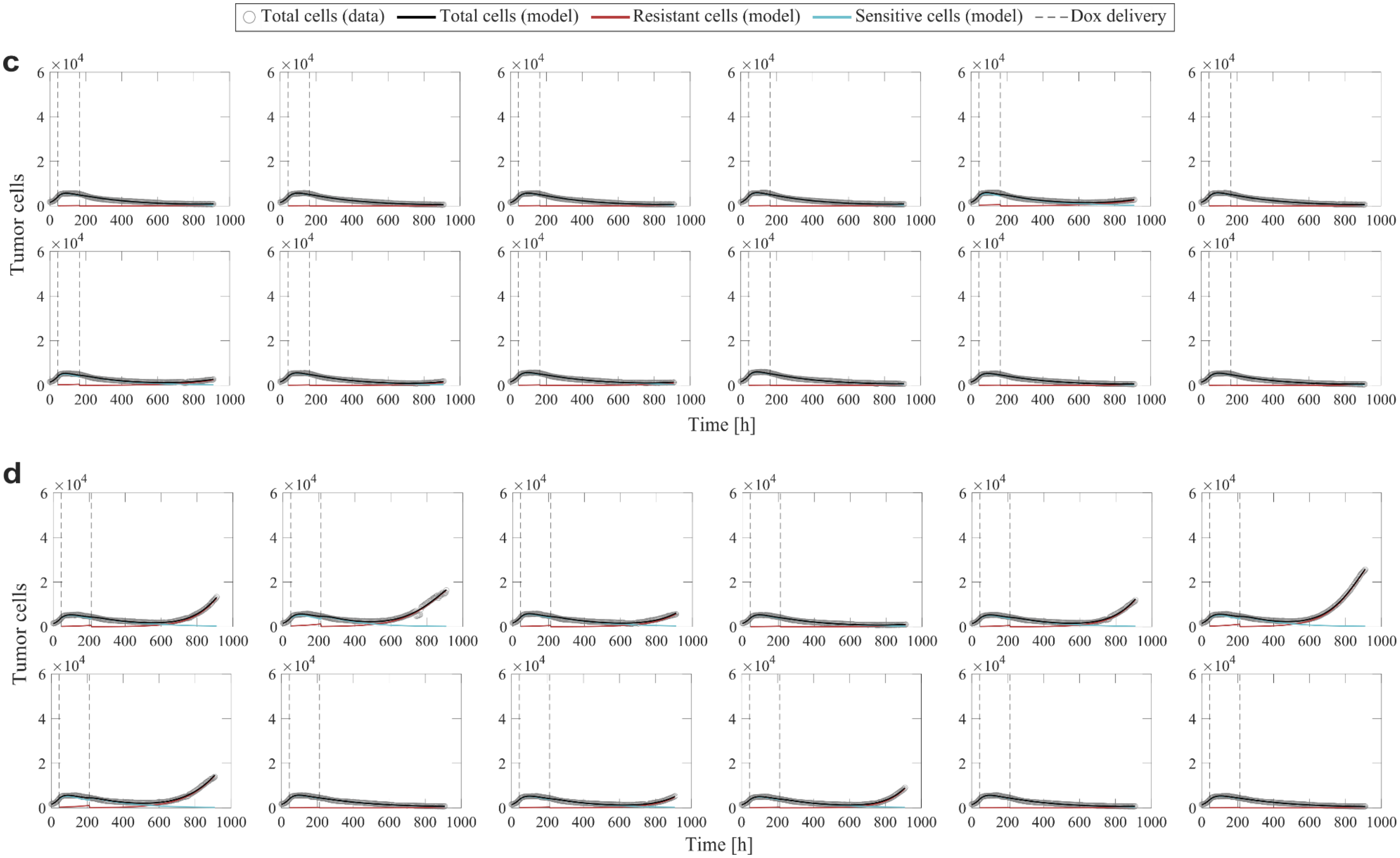

**Supplementary Figure S3 (continued).** Multiple-dose model fits for all of the time courses resulting from cells treated with two consecutive doses of 75 nM doxorubicin delivered at varying inter-treatment intervals (n = 12 for each interval; Experiment 2 in Table 1 of the main text). (**c**) 4 days; (**d**) 6 days. For (**a**)-(**d**), the multiple-dose model with constant parameters was used to fit the datasets, while for (**e**)-(**i**), the multiple-dose model with varying $f_r$ and $\gamma_d$ was used.

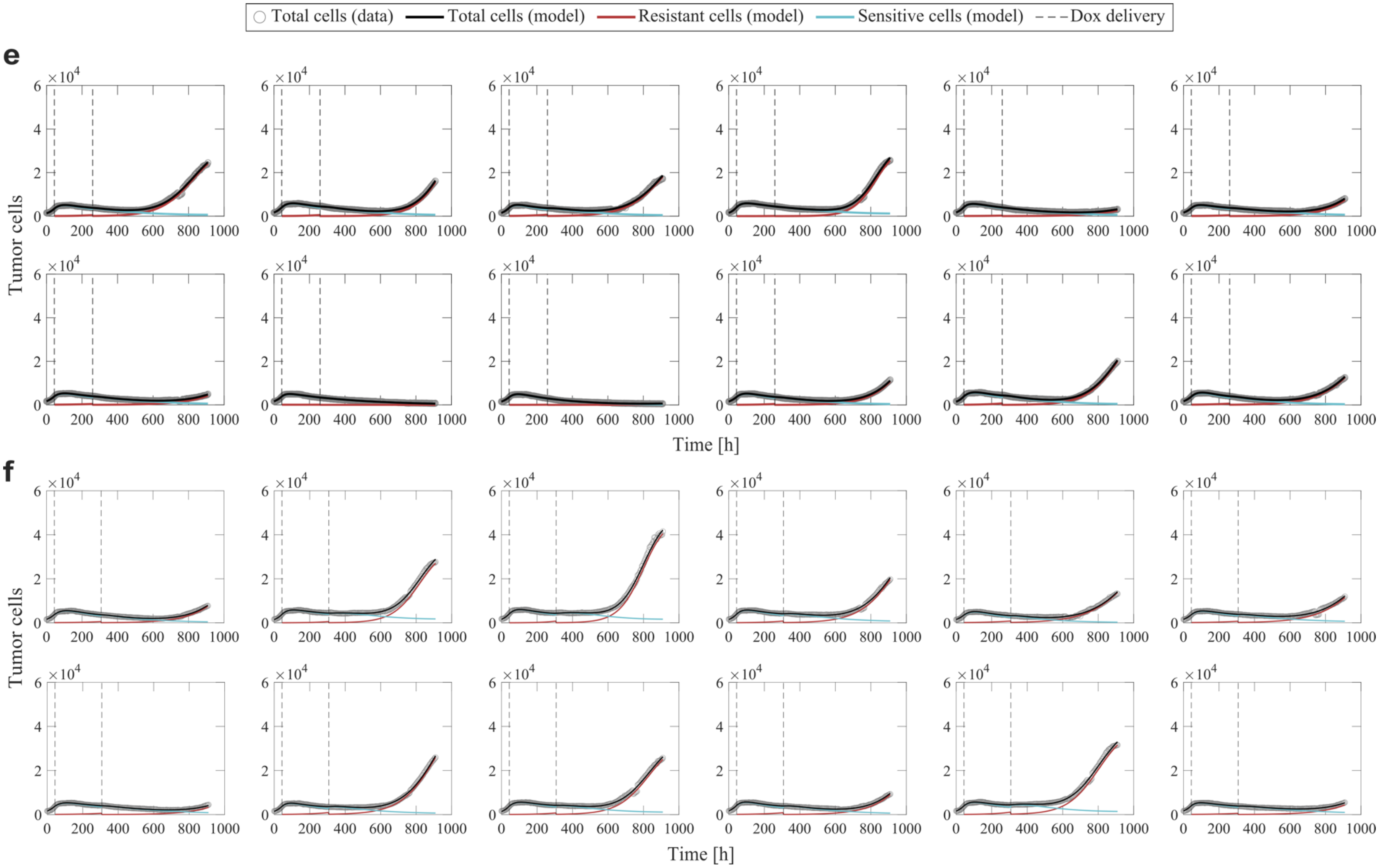

**Supplementary Figure S3 (continued).** Multiple-dose model fits for all of the time courses resulting from cells treated with two consecutive doses of 75 nM doxorubicin delivered at varying inter-treatment intervals (n = 12 for each interval; Experiment 2 in Table 1 of the main text). (**e**) 8 days; (**f**) 10 days. For (**a**)-(**d**), the multiple-dose model with constant parameters was used to fit the datasets, while for (**e**)-(**i**), the multiple-dose model with varying $f_r$ and $\gamma_d$ was used.

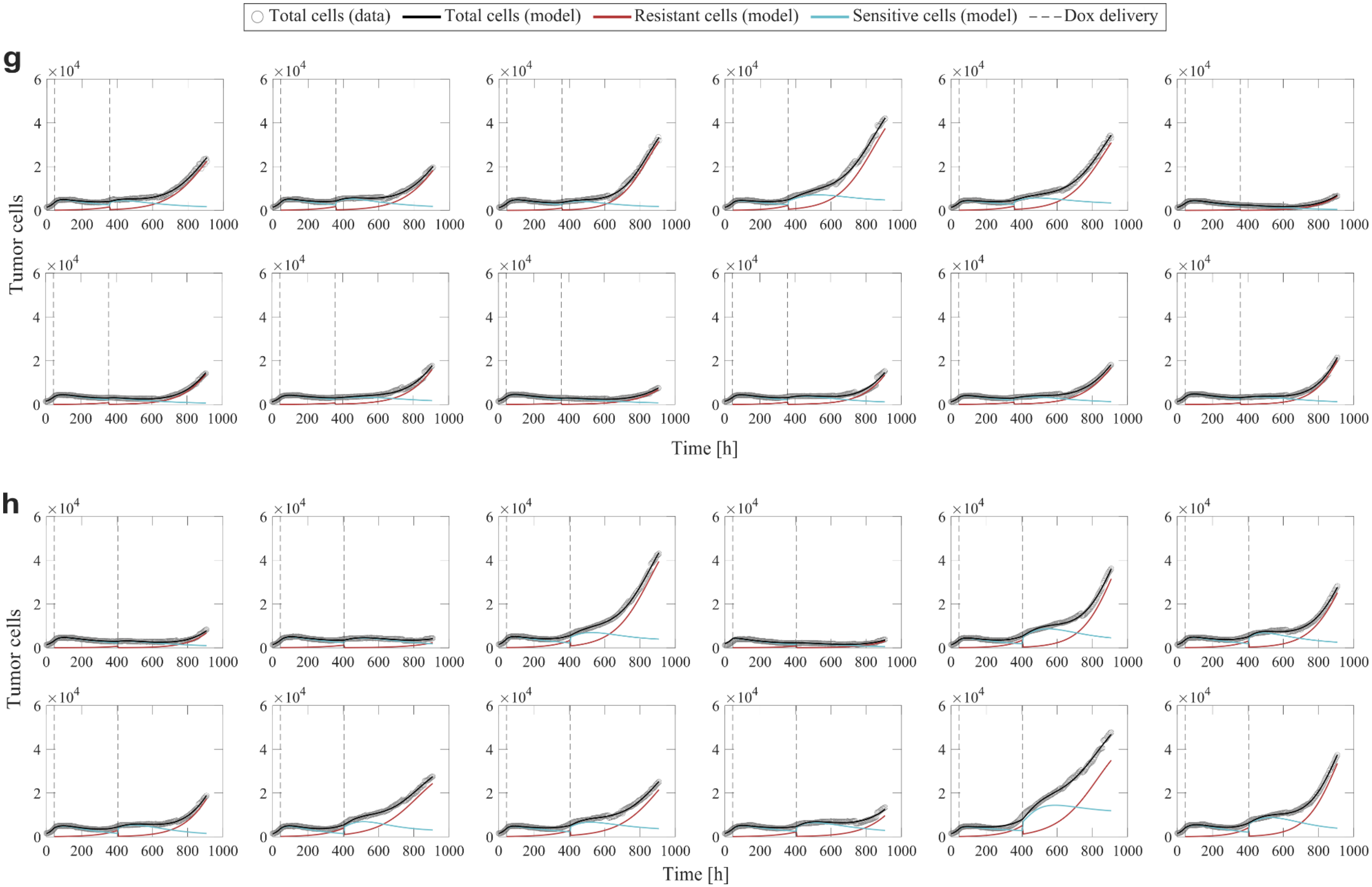

**Supplementary Figure S3 (continued).** Multiple-dose model fits for all of the time courses resulting from cells treated with two consecutive doses of 75 nM doxorubicin delivered at varying inter-treatment intervals (n=12 for each interval; Experiment 2 in Table 1 of the main text). (**g**) 12 days; (**h**) 14 days. For (**a**)-(**d**), the multiple-dose model with constant parameters was used to fit the datasets, while for (**e**)-(**i**), the multiple-dose model with varying $f_r$ and $\gamma_d$ was used.

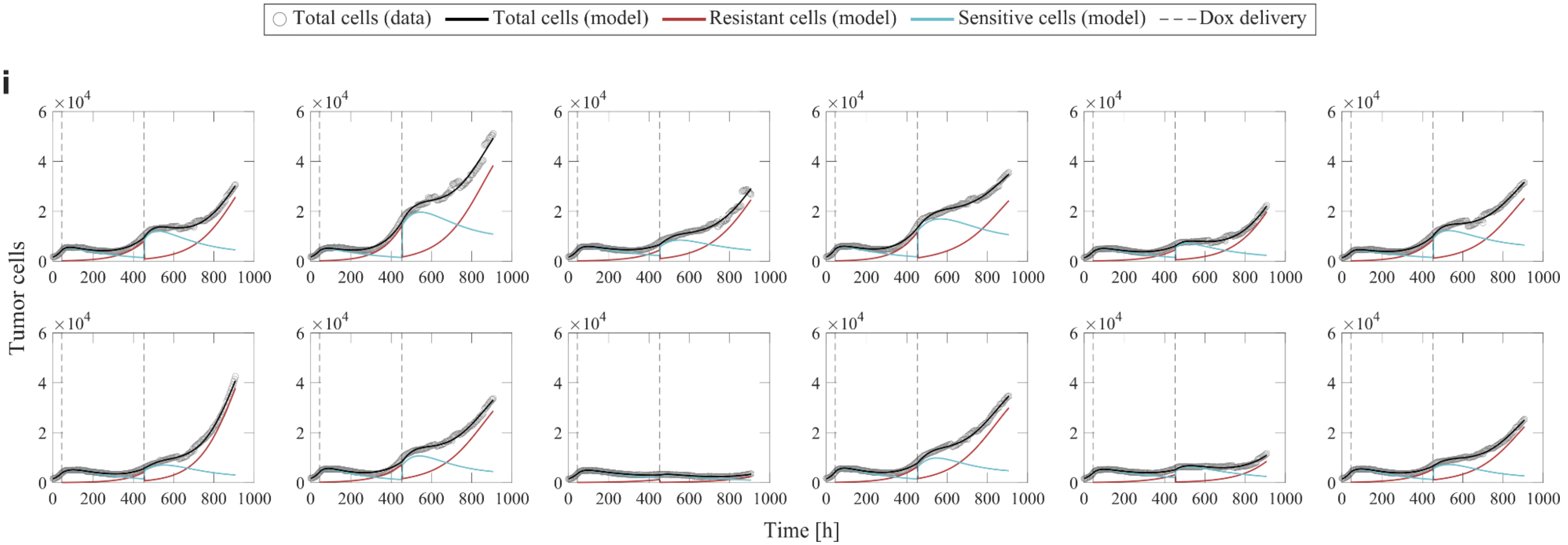

**Supplementary Figure S3 (continued).** Multiple-dose model fits for all of the time courses resulting from cells treated with two consecutive doses of 75 nM doxorubicin delivered at varying inter-treatment intervals (n=12 for each interval; Experiment 2 in Table 1 of the main text). (**i**) 16 days. For (**a**)-(**d**), the multiple-dose model with constant parameters was used to fit the datasets, while for (**e**)-(**i**), the multiple-dose model with varying $f_r$ and $\gamma_d$ was used.

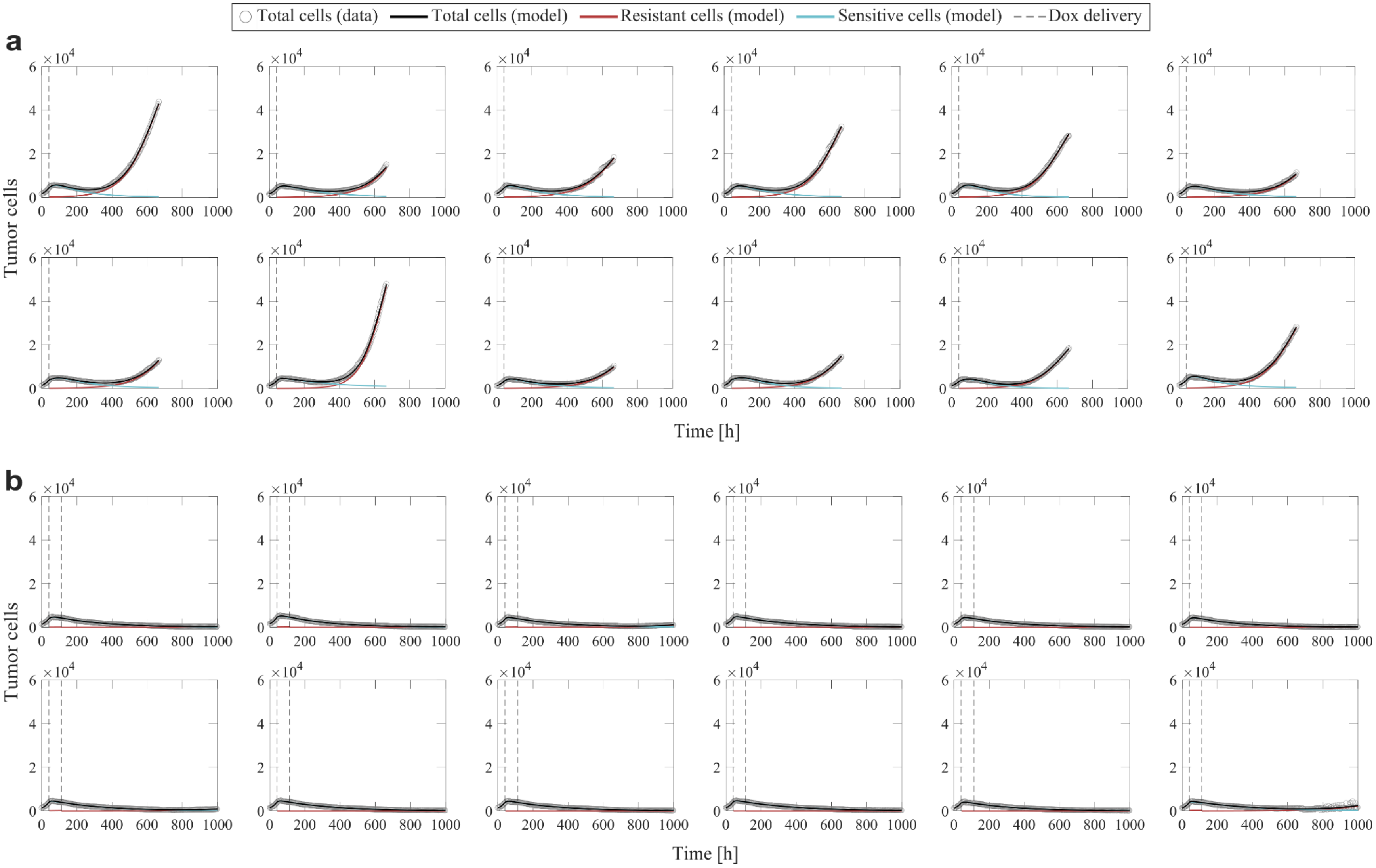

**Supplementary Figure S4.** Multiple-dose model fits for all of the time courses resulting from cells treated with a varying number of 75 nM doxorubicin doses delivered at 2-day inter-treatment intervals (n = 12 for each total dose number; Experiment 3 in Table 1 of the main text). **(a)** 1 dose; **(b)** 2 doses. The multiple-dose model with constant parameters was used to fit the datasets.

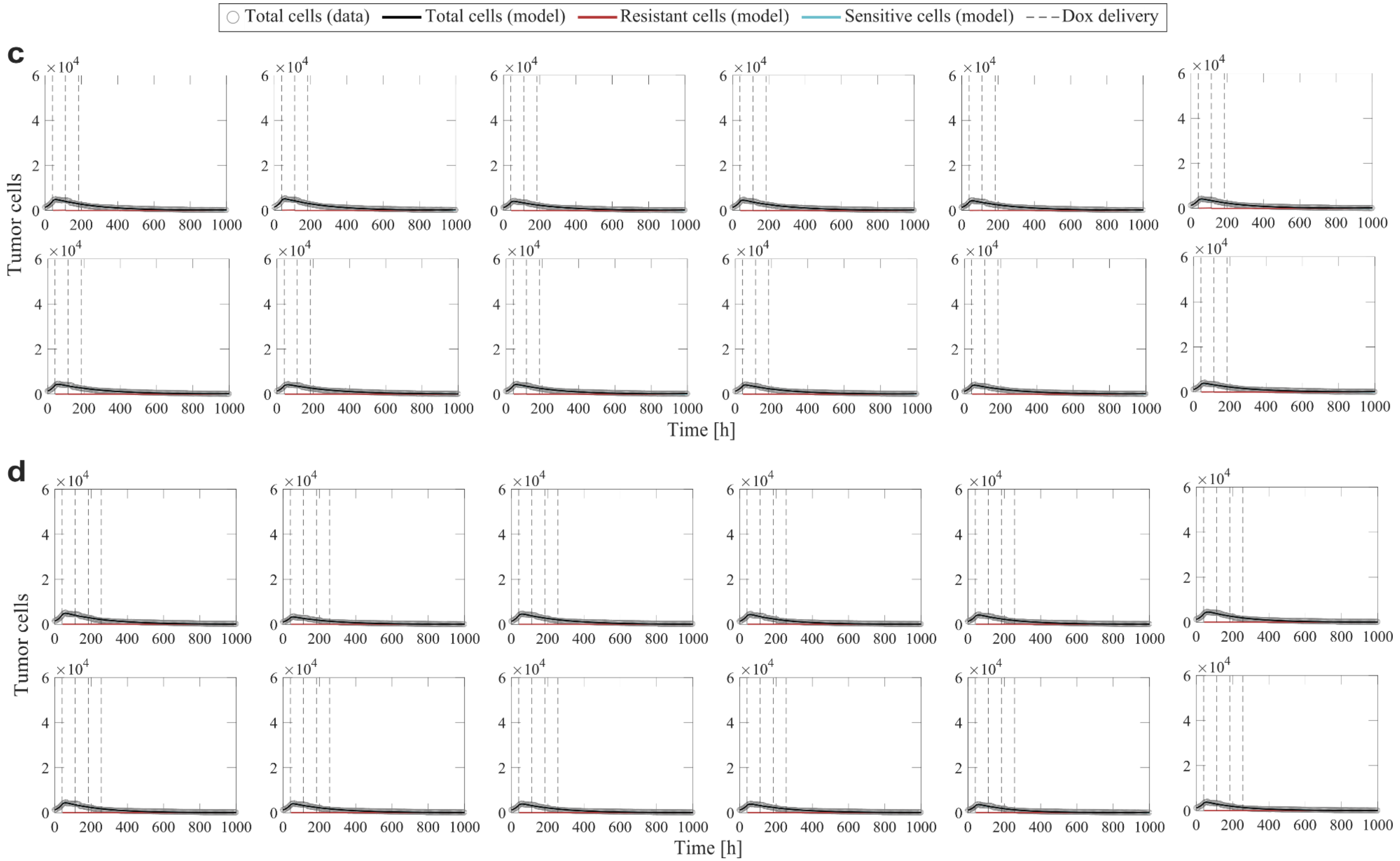

**Supplementary Figure S4 (continued).** Multiple-dose model fits for all of the time courses resulting from cells treated with a varying number of 75 nM doxorubicin doses delivered at 2-day inter-treatment intervals (n = 12 for each total dose number; Experiment 3 in Table 1 of the main text). **(c)** 3 doses; **(d)** 4 doses. The multiple-dose model with constant parameters was used to fit the datasets.

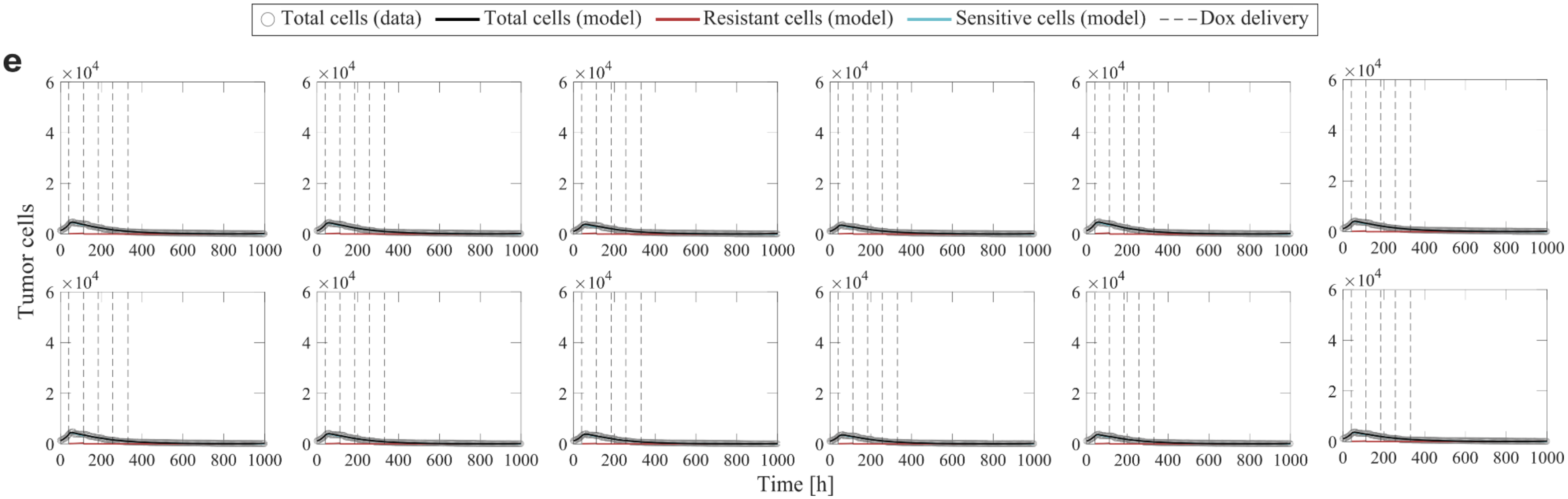

**Supplementary Figure S4 (continued).** Multiple-dose model for all of the time courses resulting from cells treated with a varying number of 75 nM doxorubicin doses delivered at 2-day inter-treatment intervals (n = 12 for each total dose number; Experiment 3 in Table 1 of the main text). (**e**) 5 doses. The multiple-dose model with constant parameters was used to fit the datasets.

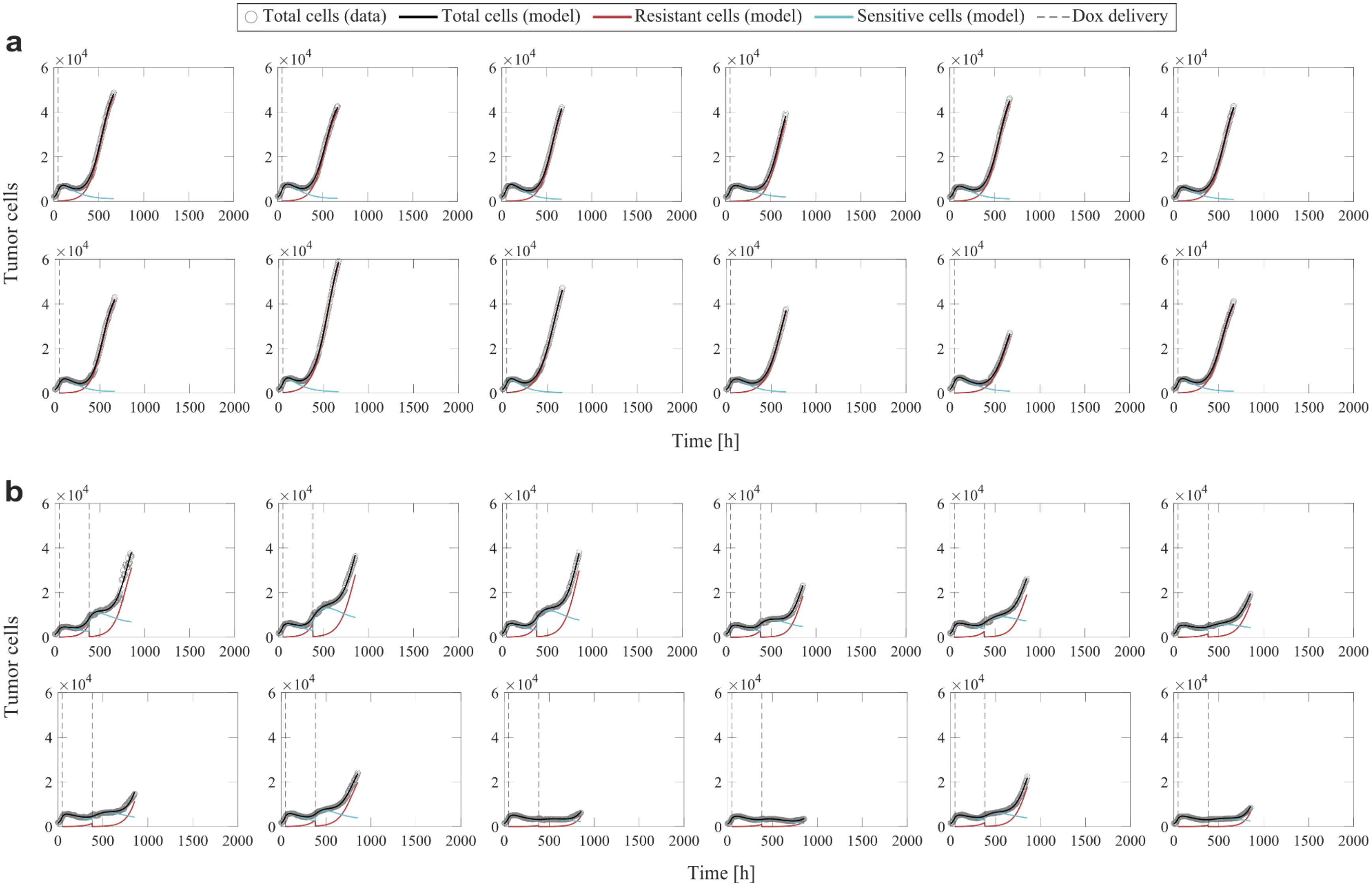

**Supplementary Figure S5.** Multiple-dose model fits for all of the time courses resulting from cells treated with a varying number of 75 nM doxorubicin doses delivered at 2-week inter-treatment intervals (n = 12 for each total dose number; Experiment 3 in Table 1 of the main text). **(a)** 1 dose; (**b**) 2 doses. The multiple-dose model with varying $f_r$ and $\gamma_d$ was used to fit the datasets.

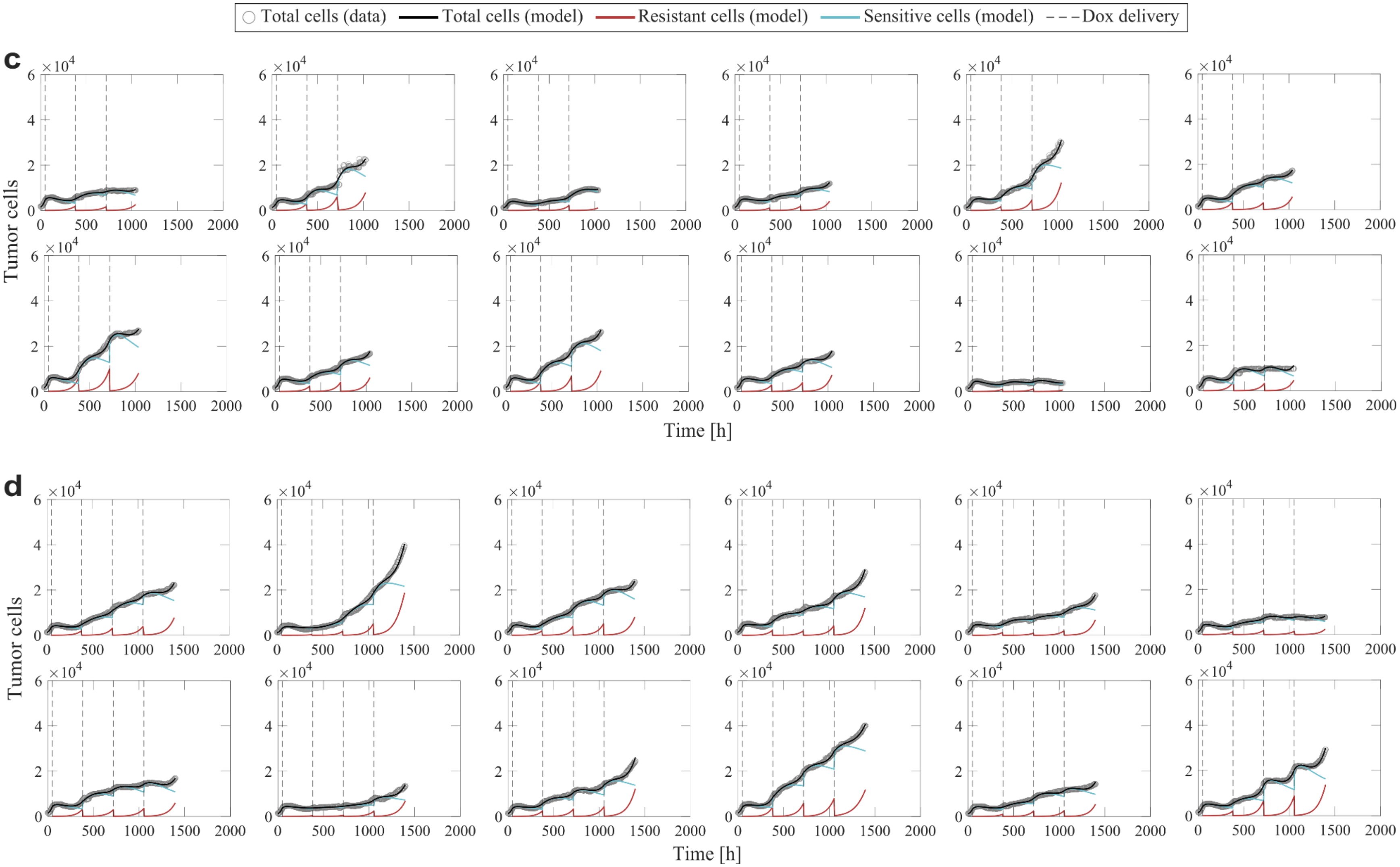

**Supplementary Figure S5 (continued).** Multiple-dose model fits for all of the time courses resulting from cells treated with a varying number of 75 nM doxorubicin doses delivered at 2-week inter-treatment intervals (n = 12 for each total dose number; Experiment 3 in Table 1 of the main text). **(c)** 3 doses; **(d)** 4 doses. The multiple-dose model with varying $f_r$ and $\gamma_d$ was used to fit the datasets.

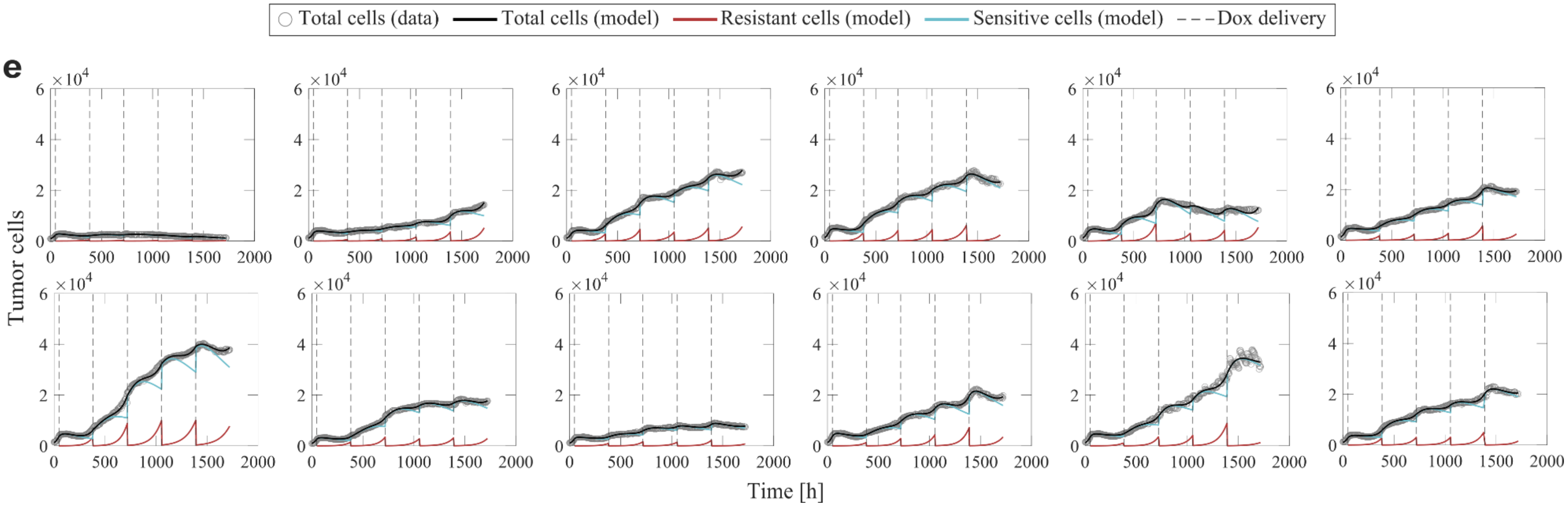

**Supplementary Figure S5 (continued).** Multiple-dose model fits for all of the time courses resulting from cells treated with a varying number of 75 nM doxorubicin doses delivered at 2-week inter-treatment intervals (n = 12 for each total dose number; Experiment 3 in Table 1 of the main text). (**e**) 5 doses. The multiple-dose model with varying $f_r$ and $\gamma_d$ was used to fit the datasets.
